# Single-molecule imaging suggests compact and spliceosome dependent organization of long introns

**DOI:** 10.1101/2021.10.27.466141

**Authors:** Srivathsan Adivarahan, A.M.S. Kalhara Abeykoon, Daniel Zenklusen

## Abstract

Intron removal from pre-mRNAs is a critical step in the processing of RNA polymerase II transcripts, required to create translation competent mRNAs. In humans, introns account for large portions of the pre-mRNA, with intronic sequences representing about 95% of most pre-mRNA. Intron length varies considerably; introns can be as short as a few to hundreds of thousands of nucleotides in length. How nascent long intronic RNA is organized during transcription to facilitate the communication between 5’ and 3’ splice-sites required for spliceosome assembly however is still poorly understood. Here, we use single-molecule fluorescent RNA in situ hybridization (smFISH) to investigate the spatial organization of co- and post-transcriptional long introns in cells. Using two long introns within the *POLA1* pre-mRNA as a model, we show that introns are packaged into compact assemblies, and when fully transcribed, are organized in a looped conformation with their ends in proximity. This organization is observed for nascent and nucleoplasmic pre-mRNAs and requires spliceosome assembly, as disruption of U2 snRNP binding results in introns with separated 5’ and 3’ ends. Moreover, interrogating the spatial organization of partially transcribed co-transcriptional *POLA1* intron 35 indicates that the 5’ splice site is maintained proximal to the 3’ splice site during transcription, supporting a model that 5’ splice site tethering to the elongating polymerase might contribute to spliceosome assembly at long introns. Together, our study reveals details of intron and pre-mRNA organization in cells and provides a tool to investigate mechanisms of splicing for long introns.

## Introduction

In eukaryotes, mRNAs are transcribed as precursor mRNAs (pre-mRNAs) that associate with RNA-binding proteins to form RNA-proteins complexes (RNPs), that undergo multiple processing steps before their export to the cytoplasm for translation. One critical step of the mRNA processing pathway is the removal of introns, a process carried out by the spliceosome, a megadalton complex comprising five U snRNPs and numerous non snRNP proteins that assemble at the 5’ splice site (ss), branchpoint site, and the 3’ ss of an intron (Will and Lührmann 2011). Splice site recognition and spliceosome assembly need to be robust as errors in splicing can result in mRNAs either quickly degraded or translated into truncated and potentially toxic proteins. Consistent with this, defects in splicing are associated with a wide range of diseases, including cancer and neurodegenerative diseases (Scotti and Swanson 2015). While spliceosome assembly and splicing catalysis have been extensively studied, little is known about how an intron is co-transcriptionally packaged and organized and how this facilitates the assembly of the spliceosome across the ends of introns, often separated by tens to hundreds of thousands of nucleotides (Wilkinson, Charenton, and Nagai 2019).

The structural and compositional organization of nascent intronic RNA occurs co-transcriptionally, likely through the binging by heterogeneous nuclear ribonucleoproteins (hnRNP) proteins and other RNA-binding proteins (Singh et al. 2015). Such a role is best described for hnRNP C that was shown to form tetramers able to package RNAs *in vitro* into compact particles of uniform size and shape, a function supported by *in vivo* CLIP experiments (Huang et al. 1994; Van Nostrand et al. 2016; König et al. 2010). In addition, RNA as a single-stranded polymer can form extensive secondary and tertiary structures, and recent transcriptome-wide chemical mapping data has identified extensive folding within intronic regions of the pre-mRNA, suggesting that RNA structure also plays an important role in intron compaction (Sun et al. 2019). However, the impact of RNA folding and RNA-binding proteins on intron compaction is largely unknown, as to date, the architecture of single introns has not been visualized in cells, and how co-transcriptional assembly of introns and its subsequent organization facilitates its splicing is yet to be determined.

In cells, splicing can occur co-transcriptionally and post-transcriptionally, with the majority of the introns known to be spliced when the RNA is still tethered to the polymerase, with co-transcriptional splicing thought to increase the efficiency and accuracy of splicing (Coulon et al. 2014; Vargas et al. 2011; Bentley 2014). However, how the spliceosome functionally pairs the ends of introns in metazoans, especially when separated by several thousands of nucleotides, and how the transcriptional machinery facilitates this process has been largely unclear. Interactions between RNA Pol II and U1 snRNP led to the model that the U1 snRNP bound to the 5’ ss could be tethered to the elongating polymerase keeping the 5’ splice site in proximity when the polymerase reaches the 3’ end, a model supported by a recent cryo-EM structure of a U1 snRNP-Pol II complex that showed a direct interaction between U1-70K of the U1 snRNP and RPB2 and RPB12 subunits of RNA Pol II (Nojima et al. 2018; Harlen et al. 2016; David et al. 2011; Robert et al. 2002; Hollander et al. 2016; Zhang et al. 2021). However, evidence for such interactions or the effect of tethering on intron organization and splicing has not yet been observed in cells.

Here, we use single-molecule resolution fluorescent RNA in situ hybridization (smFISH) in combination with structured illumination microscopy (SIM) super-resolution microscopy to determine the organization of introns, an approach we have previously used to study mRNA organization in cells (Adivarahan et al. 2018). We find that the introns are packaged co-transcriptionally into compact particles, with the organization of the intron altered during transcription. While partially transcribed introns have the 5’ end closer to the furthest transcribed region, introns containing the 3’ end are assembled with the ends in proximity. This conformation of fully transcribed introns depends on the assembly of the spliceosome, disruption of which alters intron organization, opening up the intron in RNP. Together, our results present the first overview of intron organization and compaction in cells and provide evidence to support co-transcriptional tethering of the 5’ ss to allow spliceosome assembly at long introns.

## Results

### Introns are organized as compact particles with the ends in proximity

To study the spatial organization and compaction of introns, we chose *POLA1*, encoding a subunit of DNA polymerase, as a model gene. The 303 kb long *POLA1* gene is transcribed into a pre-mRNA containing multiple long introns, with the last intron, intron 36, being 65,255 nt in length (Figure 1 A). To determine if POLA1 pre-mRNAs can be detected in cells, we hybridized smFISH probes to a 5’ region of intron 36 and the 5’ exonic region in paraformaldehyde-fixed HEK293T cells. As expected, 5’ exonic signals are detected in the nucleus and cytoplasm, with nuclear signals clustering in one or two sites representing sites of nascent POLA1 pre-mRNA transcription (Figure 1 B). Intron signals are observed at the site of transcription and in the nucleoplasm, often close to transcription sites. Whereas signal intensity of nucleoplasmic introns is largely uniform, intron signals at the site of transcription vary, indicating the presence of pre-mRNAs containing one or more nascent introns. Most intronic signals in the nucleoplasm co-localize with exonic signals, suggesting that these spots represent pre-mRNAs that have not been spliced co-transcriptionally, whereas intron signals not co-localizing with exonic signals represent intron lariats. Quantification of nucleoplasmic intron signals showed 185 out of 209 individual spatially distinct intron signals co-localizing with exon signal, indicating that most single nucleoplasmic introns represent pre-mRNAs. Overall, we detect single introns at sites of transcription, as well as non-spliced nucleoplasmic pre-mRNAs and nucleoplasmic intron lariats. Henceforth, we used this approach to study intron organization.

**Figure 1:**
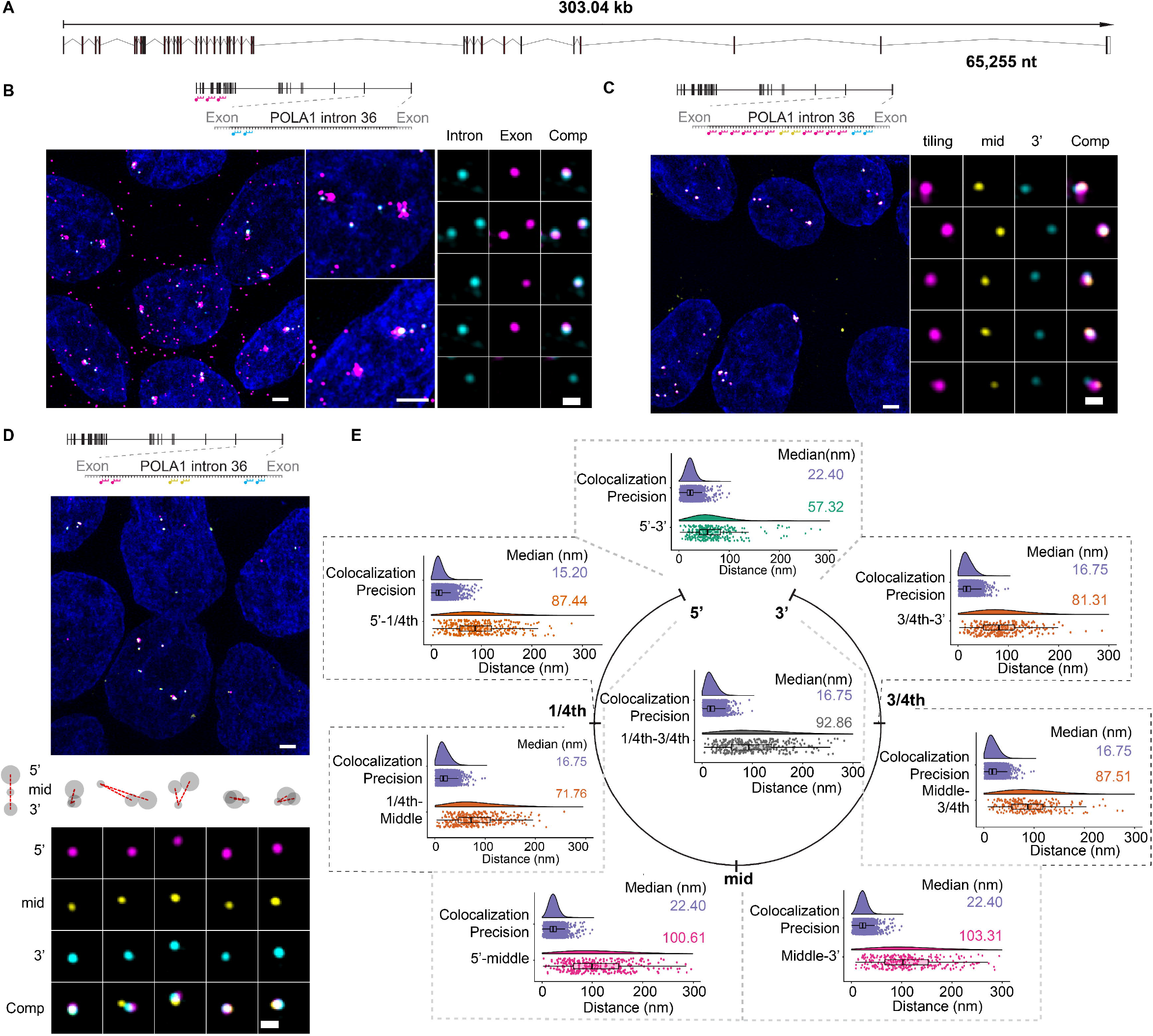
Organization of intron 36 of the POLA1 gene. **(A)** Exon-intron structure of human POLA1 gene for the transcript ENST00000379059, with the length of intron 36 shown at the bottom, smFISH images using probes hybridizing to **(B)** 5’ exons (magenta) and 5’ region of the intron 36 (cyan) (Probe Set#3, Table S 2) in paraformaldehyde-fixed HEK293T cells **(C)** the middle (yellow), 3’ (cyan), and tiling along the intron (magenta) (Probes Set#7, Table S 2) **(D)** 5’(magenta), middle (yellow), 3’ (cyan) regions (Probes Set#6, Table S 2) Nuclei are visualized by DAPI staining (blue). Magnified images of individual RNAs are shown at the bottom or on the right, and cartoons depicting different RNA conformations are shown below the images (D). Schematic position of probes shown on top. **(E)** Raincloud plots for distances between different regions. Individual plots show distance distribution of co-localization precision, distances for POLA1 introns as violin plots. The box plot shows the first quartile, median and third quartile and the distances corresponding to single RNAs are shown as spots overlayed on top of the box plots. Median distances are shown on the right. Scale bars, 2 µm in larger and zoomed-in images, and 500 nm in images showing single introns.

Next, we determined how this 65,255 nt long intron is spatially organized. We first hybridized cells with a combination of probes tiling along the length of the intron (2 probes ∼ every 1,000 nt) and probes targeting the middle and 3’ regions of the intron (Figure 1 C). Previously, studying the spatial organization of cytoplasmic *MDN1* mRNAs (18,413nt) using tiling probes showed an elongated organization with an irregular shape and a volume greater than the diffraction limit (Adivarahan et al. 2018). Signal for probes tiling intron 36 revealed a nearly circular shape near the diffraction limit for most introns, indicating a highly compact and possibly globular molecule organization (Figure 1 C). Signals to the middle and 3’ regions on intron 36 overlapped with the tiling signal. However, despite the small size of the tiling spots, we observed a spatial separation of middle and 3’ regions within this compact particle.

To better understand the organization of this intron, we replaced the tiling probes with probes targeting the 5’ region of the intron, allowing us to quantify the relative position of the three regions with respect to each other. We measured the distances between different regions by first localizing the center of the signal using 3D Gaussian fitting, converting the 3D coordinates to 2D coordinates, and measuring the distances between co-localizing signals from different channels. As a measure of our experimental and localization error, we targeted probes to a ∼1,200 nt region of 18,413 nt-long *MDN1* mRNA alternately labelled with Cy3 and Cy5 and measured the distances between signals from the two channels to obtain what we term as “Co-localization precision”, typically ranging between 15-25 nm. Figure 1 D shows introns imaged with the probes against the 5’, middle and 3’ regions, each region separated by ∼30,000nt of RNA (Supplementary Figure 1). These measurements showed that though the introns had compact conformations, the introns were almost always organized with the ends in proximity compared to the middle region. Measuring distances between different regions result in end-to-end distances with a median of ∼57 nm compared to distances of ∼100 nm between the ends and the middle region of the intron (Figure 1 E).

To further dissect the size and organization of this intron, we divided the intron into four sections and designed smFISH probes against the boundaries of these sections (5’, 1/4^th^, mid, 3/4^th^ and 3’, see Supplementary Figure 1), which were then used in various combinations, allowing us to calculate the distances between these regions (Table S 2). These measurements show that intron 36 is organized into particles with distances that are longest between the ends of the intron and the middle region and the shortest between the two ends of the intron. All other regions were separated by end-to-end and end-to-middle distances, suggesting that intron 36 molecules are possibly organized in a looped conformation (Figure 1 E and Supplementary Figure 2). Furthermore, the distance measurements suggest the intron particles have a diameter of ∼100 nm. Together, our results show that we can detect individual POLA1 intron 36 molecules, most of which are still part of pre-mRNAs. Furthermore, a large fraction of POLA1 intron 36 molecules are found near transcription sites, likely representing pre-mRNAs that are spliced post-transcriptionally and organized into compact particles with looped conformations having their ends in proximity.

### Organization of fully and partially transcribed nascent POLA1 introns 35

Next, we wanted to test whether other introns show similar organization as observed for intron 36. We chose the 42,330nt long intron 35 of the POLA1 gene and hybridized probes against the 5’ end of the intron and the 5’ end of POLA1 exons (Figure 2 A). Compared to intron 36, which is found as part of nucleoplasmic pre-mRNAs and at the site of transcription, intron 35 smFISH signals were mostly observed at sites of transcription (Figure 2 B), suggesting more efficient co-transcriptional splicing of this intron and a fast turnover of the lariat. Combined with the low transcription frequency of the POLA1 gene in HEK293T cells, we found that this allowed us to identify single introns at the transcription site (Figure 2 B).

**Figure 2:**
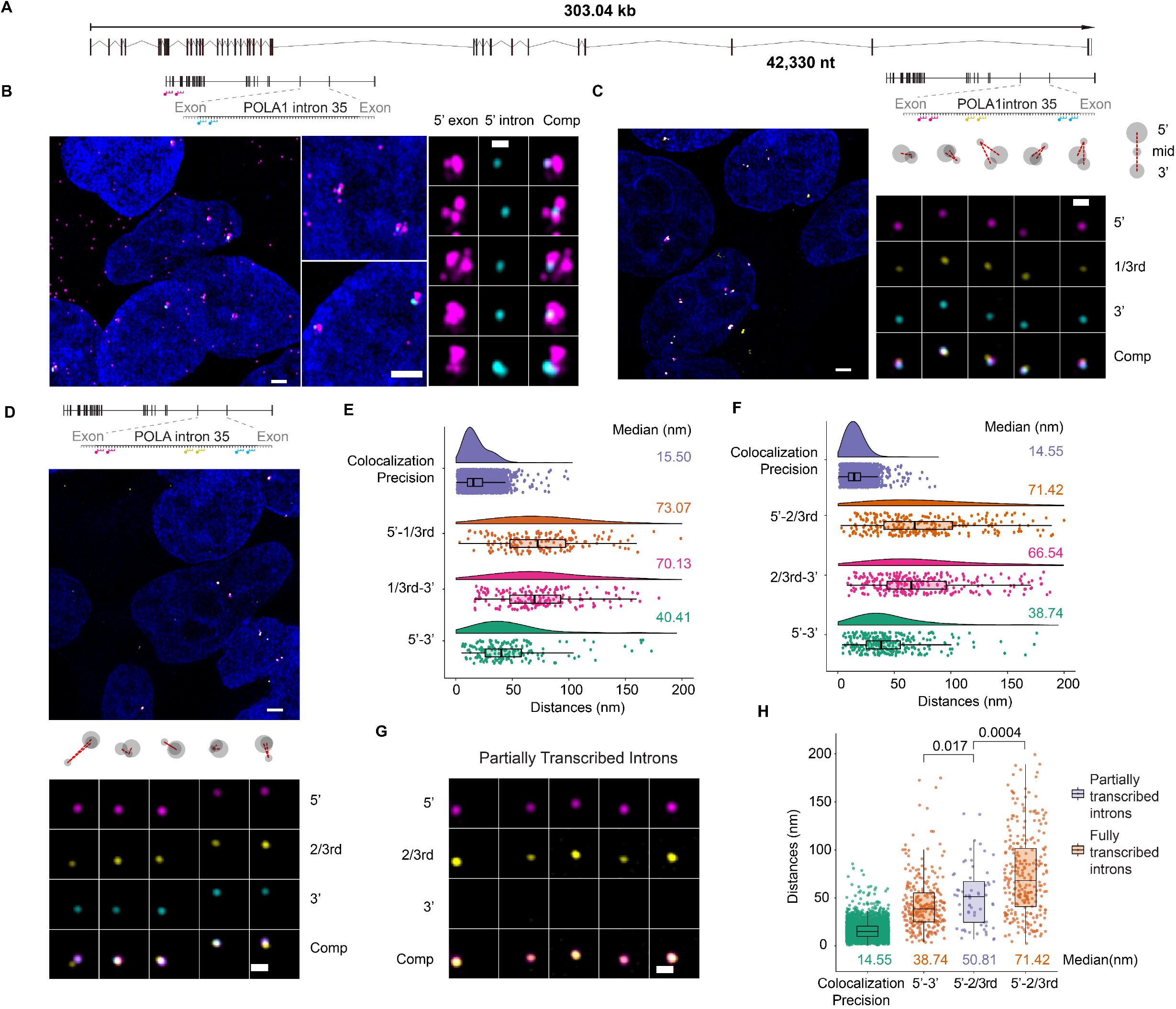
Organization of partially and fully transcribed intron 35 molecules. **(A)** Exon-intron structure of human POLA1 gene for the transcript ENST00000379059, with the length of intron 35 shown at the bottom **(B)** smFISH images using probes hybridizing to 5’ exons (magenta) and 5’ end of the intron 36 (cyan) (Probe Set#4, Table S 2) in paraformaldehyde-fixed HEK293T cells. **(C, D)** smFISH using probes hybridizing to 5’(magenta), 3’ (cyan) regions and either 1/3^rd^ or 2/3^rd^regions (yellow) (Probes Set#13 and #14 respectively, Table S 2) and cartoons depicting different RNA conformations. Nuclei are visualized by DAPI staining (blue). Magnified images of individual RNAs are shown at the bottom or the right. Schematic position of probes shown on top. **(E, F)** Raincloud plots for distances between different regions. Individual plots show distance distribution of co-localization precision, distances for POLA1 intron 35 shown as violin plots. The box plot shows the first quartile, median and third quartile and the distances corresponding to single RNAs are shown as spots overlayed on top of the box plots. Median distances are shown on the right, **(G)** Magnified smFISH images of partially transcribed individual RNAs using smFISH probes hybridizing to the 5’(magenta), 2/3^rd^ (yellow) and 3’ regions (cyan) (Probes Set#14, Table S 2). **(H)** Box plot with distances between different regions for partially and wholly transcribed introns 35 from (E) and (G). The box plot shows the first quartile, median and third quartile and the individual RNAs shown as spots overlayed on top of the box plots. P-values calculated using Kolmogorov–Smirnov test are shown above. Scale bars, 2 µm in larger and zoomed in images, and 500 nm in images showing single introns or transcription sites in (B), (C), (E) and (G).

To get a detailed understanding of this intron’s organization, we divided the intron into three regions and designed probes against the boundaries of these regions (Supplementary Figure 1). Three-colour imaging of the intron using probes hybridizing against the 5’, 3’ ends with the third region reveals an organization of intron 35 to be similar to what was observed for intron 36, with the median end-to-end distances ∼40 nm and the distances between the ends and the third region larger than the end-to-end distances (Figure 2 C-F).

As most intron 35 molecules were observed at the site of transcription, it also allowed us to visualize the organization of partially transcribed introns. If the co-transcriptional assembly of introns is involved in ensuring the proximity of their ends, as described by the U1-snRNP-Pol II tethering model, we would expect intron organization to be altered during transcription. To test this model, we quantified distances for introns that contained signals for the 5’ and the 2/3^rd^ region of the POLA1 intron 35 but were missing the 3’ signal, indicative of transcription not yet reaching the 3’ end. We argued that if the 5’ ss stays attached to the elongating polymerase complex, partially transcribed introns containing signals against just the 5’ and 2/3^rd^ regions should have polymerases closer to the 2/3^rd^ region, resulting in shorter distances than observed for the same regions within introns containing the 3’ signal. Visualizing single introns showed an increased overlap of the 5’ and 2/3^rd^ signals when the 3’ signal was absent, which was reflected in shorter end-to-end distances (Figure 2 G-H). Figure 2 H shows that 5’-2/3^rd^ distances for partially transcribed introns lie between the end-to-end and 5’-2/3^rd^ distances for fully assembled introns. Together, our observations show that POLA1 intron 35 has an organization similar to intron 36, with a looped conformation, which is altered during transcription in a manner consistent with a model that suggests 5’ ss tethering to an elongating polymerase.

### Spliceosome assembly modulates intron organization of nascent and nucleoplasmic pre-mRNAs

The intronic organization with the 5’ and 3’ ends in proximity is expected for introns in the process of being spliced as well as intron lariats, but not for nucleoplasmic pre-mRNAs, except if these pre-mRNAs would contain, at least, partially assembled spliceosome complexes or if the co-transcriptional assembly of introns was, by itself, sufficient for maintaining this proximity. To distinguish between these possibilities, we inhibited splicing with Pladienolide B (PB), a small molecule that binds to SF3B1 and interferes with U2 snRNP assembly at the branch point. Consistent with splicing inhibition and transcript release from the site of transcription without splicing, we find an increased number of intron spots for both *POLA1* introns (introns 35 and 36) in the nucleoplasm upon treatment with 100 nM PB for four hours (Supplementary Figure 3). Treatment with PB also resulted in an increase in nuclear (pre-) mRNAs containing just the 5’ exon signal without the 3’ exon signal outside the site of transcription, consistent with an increased frequency of premature termination, as described earlier (Sousa-Luís et al. 2021) (Supplementary Figure 6 A).

To determine if splicing inhibition altered intronic conformations, we targeted probes to the 5’ and 3’ ends of both introns in combination with probes targeting the third region of each intron (middle in case of intron 36 and 1/3^rd^ and 2/3^rd^ in case of intron 35). When all three signals were visualized for these introns, we noticed an increased separation of signals corresponding to the 5’ and 3’ ends of introns in cells treated with PB (Figure 3 A and Supplementary Figure 4 A, B). This increased separation was reflected in our distance measurements where we see the median end-to-end distances increases from ∼57 nm to ∼144 nm for intron 36 and from ∼40 nm to > 120 nm for intron 35, suggesting that establishment of the proximity between the 5’ and 3’ ends of introns is dependent on the assembly of U2 snRNP (Figure 3 B, C, F and Supplementary Figure 4 C-E). Interestingly, we also observe a small increase in distances between the ends and the third region for both introns. This might suggest an overall increased decompaction of the intron, possibly due to a partial unfolding of an otherwise compact intron (Figure 3 C, F and Supplementary Figure 4 C-E). Furthermore, analyzing distances of individual introns shows that treatment with PB results in introns with end-to-end distances larger than distances between ends and the third region and an increase in the angle formed between the 5’, third region and 3’ regions, suggesting an overall change in the organization of introns upon inhibition of U2 assembly (Figure 3 D, E, G, H and Supplementary Figure 4 F-G).

**Figure 3:**
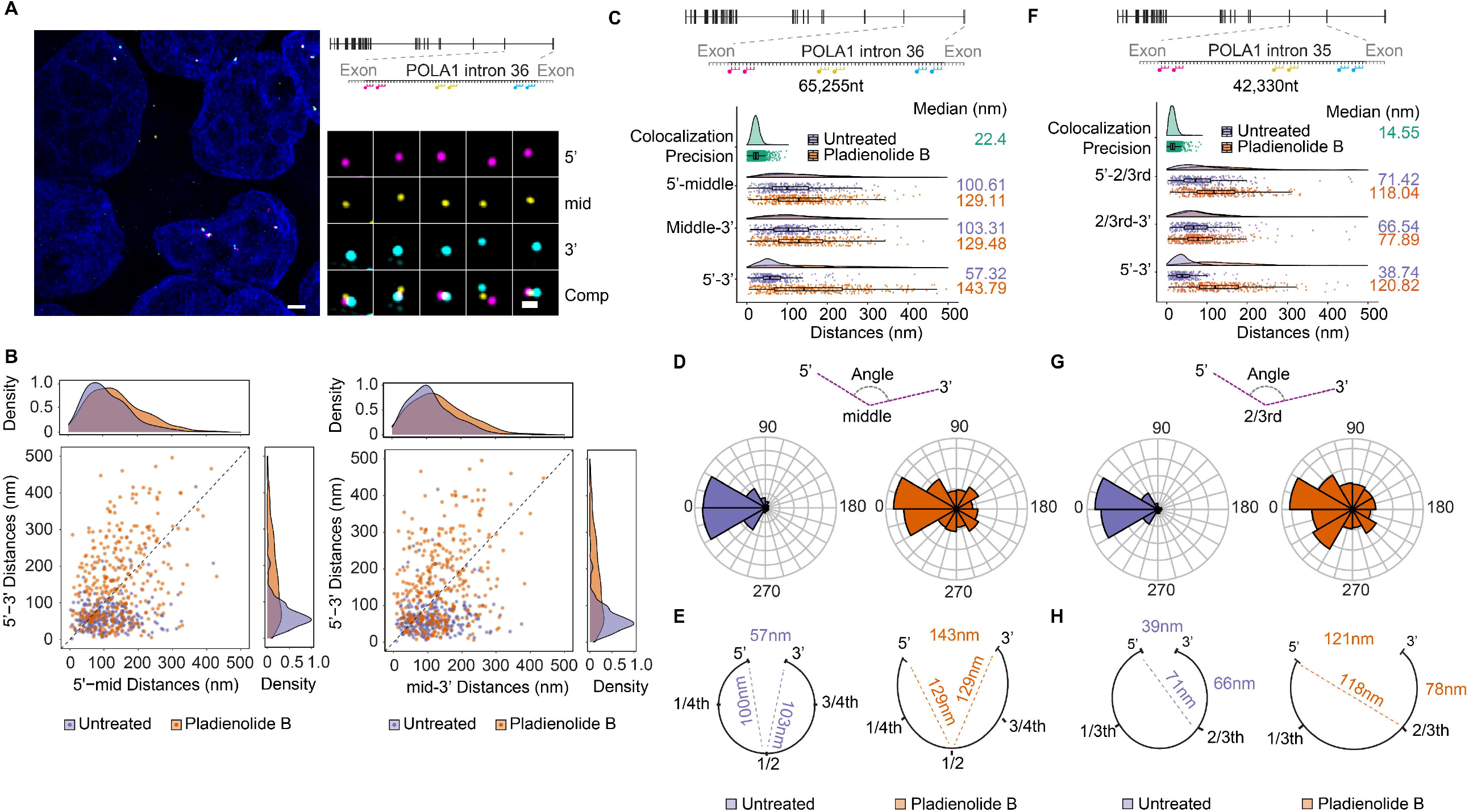
Intron Organisation is altered upon inhibition of U2 assembly. **(A)** smFISH images in HEK293T cells treated with 100 nM Pladienolide B for 4h. Nuclei are visualized by DAPI staining (blue). Magnified images of individual RNAs are shown on the right. Schematic position of probes shown on top. Probes hybridize to 5’(magenta), middle (yellow), 3’ (cyan) regions of the intron 36 (Probes Set#6, Table S 2), **(B)** Scatter plot showing 5’mid and mid-3’ distances for individual introns for intron 36 for mock and Pladienolide B treated HEK293T cells). Frequency distribution is shown on top and on the right. **(C, F)** Raincloud plots for distances between different regions for untreated and treated cells. Individual plots show distance distribution of co-localization precision and distances for POLA1 introns as violin plots. The box plot shows the first quartile, median and third quartile and the individual RNAs shown as spots. Median distances are shown on the right. **(D, G)** Angle histogram plot showing angle formed between the three regions of intron 36 and 35 as represented in the cartoon. **(E, H)** Cartoon representation of median distances between different regions of intron 36 and 35 as observed in C, F. Scale bars, 2 µm in larger images, and 500 nm in zoomed-in images of individual introns.

Lastly, to monitor how the changes in intronic conformations affected the overall conformations of pre-mRNPs, we hybridized probes to different exonic regions of three mRNAs - *POLA1, MDN1* and *AHNAK*. While *POLA1* and *MDN1* contain multiple introns that have previously been observed to be affected by the PB treatment, *AHNAK* is an mRNA with five exons and four introns, with the last exon having a length of 18,173nt (Kim Guisbert, Mossiah, and Guisbert 2020). Our probes to *AHNAK* were designed to hybridize to different regions within this long exon, and any changes in intronic conformations due to splicing inhibition by PB is likely not to affect the organization of the long exon within the *AHNAK* mRNA (Supplementary Figure 1). We imaged these mRNAs in untreated cells and cells treated with PB and found an increased separation between different regions for *MDN1* and *POLA1* mRNAs but not *AHNAK* mRNAs (Supplementary Figure 5 A and Supplementary Figure 6 A, D). This was reflected in our distance measurement, where the end-to-end distances for POLA1 and distances between different regions for *MDN1* increased upon PB treatment, while they were largely unaltered for *AHNAK* mRNAs (Supplementary Figure 5 C and Supplementary Figure 6 B, C, F). For three colour measurements of *AHNAK* and *MDN1*, we found that these changes in distances also resulted in changes in the size of the pre-mRNP as determined by the radius of gyration (Supplementary Figure 5 B and Supplementary Figure 6 E), indicating that intron assembly during transcription determines the final organization of pre-mRNPs. Together, our observations suggest that intron organization is dependent on the binding of U2 snRNP and possibly other spliceosomal components, inhibition of which separates the ends of the intron and alters the pre-mRNP conformation overall.

## Discussion

Although spliceosome assembly and the catalytic steps of splicing have been extensively studied, how introns are co-transcriptionally packaged and organized to facilitate the communication between the 5’ and 3’ ends are still poorly understood. Using single-molecule localization microscopy, we find that introns exist as compact particles, organized with their ends in proximity. This organization is dependent on the assembly of the spliceosome, as inhibition of branch point recognition and U2 snRNP assembly on the intron using Pladienolide B results in the separation of the introns 5’ and 3’ends at nascent, as well as nucleoplasmic pre-mRNAs. Finally, we find that intron organization is altered during transcription, with the separation between distal regions of POLA1 intron 35 changing with the length to which the intron has been transcribed, indicating that tethering of the 5’ ss to Pol II could contribute to facilitating ends of long introns to come together and initiate the assembly of a spliceosome.

### Introns are compact assemblies *in vivo*

Our previous observations showed that nuclear mRNPs are assembled into compact particles, possibly due to their linear co-transcriptional packaging, and single-molecule imaging of introns in this study shows that similar conclusions can be made for introns (Adivarahan et al. 2018). By targeting probes to multiple regions of introns 35 and 36, we determined the spatial separation between these regions and estimated these introns’ overall size within pre-mRNPs (Figure 1 and Figure 2). Our calculations suggest that intron 36 molecules containing 65,255 nt of RNA are compacted into particles with an average diameter of ∼100 nm. Similarly, intron 35 particles, containing 42,330nt of RNA, have a diameter of ∼80nm on average. Considering a hypothetical fully extended, linear mRNA with an inter-nucleotide spacing of 0.59 nm and defining compaction as the ratio of the distance between the ends of a fully extended RNA to the spherical volume of the assembled RNP, calculated using the radii, this would result in compaction of 0.073 fold for intron 36 and 0.093 fold for intron 35 (Liphardt et al. 2001). In comparison, the 80S ribosomes, one of the most extensively compacted RNPs in cells, has a diameter of ∼30nm and contain 7,216 nt of RNA, resulting in fold compaction of 0.3. High levels of compaction are also observed for viral RNPs, that require packaging into capsids with a fixed internal diameter. Performing a similar analysis on viral RNAs, we find that Hepatitis A (∼7,500 nt genomes and 24 nm inner capsid diameter), Zika viruses (∼11,000 nt and 30 nm inner capsid diameter), and SARS-COV-2 (∼30,000nt genome and ∼100nm outer capsid diameter) have compaction of 0.6, 0.4 and 0.034 respectively (Wang et al. 2017; Sirohi et al. 2016; Ke et al. 2020). While high compaction seems necessary and conceivable for these RNPs, the same is not true for mRNPs, which *in vivo* have been known to have varied compaction depending on their translational state and cellular localization (Adivarahan et al. 2018; Khong and Parker 2018). Our previous study found that translating MDN1 mRNPs (18,413 nt) to have a much more open conformation and a compaction of ∼0.006 fold; however, when translation is inhibited through the removal of ribosomes, this results in particles with average compaction of 0.13. These calculations, overall, indicate that introns are co-transcriptionally packaged into particles with a compaction level similar to the most compact state of mRNPs observed in cells – the non-translational state, but possibly less than some of the most compact RNPs found *in vivo*, such as the ribosome.

While the compaction of ribosomal RNPs starts co-transcriptionally through local RNA folding assisted by the binding of RNA binding proteins, the process for compaction of pre-mRNPs and introns, in particular, is much less understood. Similar to observations made for ribosomal subunits and several RNAs *in vitro*, RNA folding likely plays a critical role in this process (Gopal et al. 2012; Borodavka et al. 2016; Russell et al. 2002). In agreement with this, recent transcriptome-wide RNA structure analysis data has found extensive RNA folding within the intronic regions of pre-mRNPs (Sun et al. 2019). However, unlike ribosomes, the diversity in the primary sequence of introns could result in intronic RNAs forming a plethora of different thermodynamically stable structures and whether this is the cause for these RNAs having compaction lower than ribosomes is unclear.

In addition to RNA folding, compaction of POLA1 intron 35 and 36, and introns in general, is also likely achieved through packaging by RBPs assembled co-transcriptionally onto pre-mRNAs. Members of the hnRNP protein family, particularly hnRNP C, have been proposed to play a significant role in this process, and many of them are known to preferentially bind to intronic regions (Van Nostrand et al. 2016; König et al. 2010). Furthermore, hnRNP C and members of the hnRNP A/B family have been shown to form complex assemblies with RNAs, both *in vitro* and *in vivo*, packaging 700 nt of RNA into particles ∼26 nm in diameter (Huang et al. 1994; Conway et al. 1988). While the abundance of these assemblies remains unclear *in vivo*, CLIP-Seq studies have found regularly spaced peaks of hnRNP C within several introns, including intron 35 and 36 of *POLA1*, which may represent individual hnRNP C tetramers that could nucleate the formation of 40S hnRNP particles (Huang et al. 1994; König et al. 2010; Van Nostrand et al. 2016). When spread over the entire intronic region, these assemblies could help package the intronic RNA and result in the compaction that we observe for introns 35 and 36 of POLA1. Future work studying the depletion of hnRNP proteins on the intronic organization could shed some insight into this process.

Our results also show that POLA1 introns 35 and 36, though generally organized in a looped conformation, undergo an architectural change upon inhibition of U2 snRNP assembly, indicative of the opening of the loop. This opening of the loop and the resultant conformations, where the end-to-end distances are longer in comparison to distances between other regions within the introns, suggests that introns 35 and 36 are possibly packaged and compacted co-transcriptionally in a linear manner, similar to our previous observations for nuclear *MDN1* mRNAs and observations made from other studies for nuclear mRNPs (Adivarahan et al. 2018; Metkar 2018; Mehlin, Daneholt, and Skoglund 1995).

### Intron organization and spliceosome assembly

The distance measurements for introns 35 and 36 suggest that these introns have a looped organization with the ends in proximity. A possible explanation for such a conformation is the assembly of the spliceosome across the ends of the intron. Consistent with this, Pladienolide B treatment and inhibition of binding of U2 snRNP separates the ends of introns 35 and 36 while the conformations of long exons are largely unaffected. Our data, however, does not allow us to determine whether the binding of U2 snRNP, in normal conditions, results in the immediate recruitment of other spliceosomal factors and the assembly of the entire spliceosome complex. Our end-to-end measurements for POLA1 intron 35 yield a median distance of ∼40 nm, similar in size to the spliceosome complex (∼25-30 nm), distances that would be consistent with an assembled spliceosome at these introns. End-to-end distances are slightly longer for intron 36 (median ∼57nm); however, this might be in part due to the increased spread of the smFISH probes hybridizing to the 5’ or the intron (3,850nt for intron 36 vs 1,892nt for intron 35). Importantly, the looped organization of both introns is dependent on the U2 snRNP association, suggesting that assembly of U2 snRNP is critical towards keeping the ends in proximity.

Our single-molecule analysis also shows that a large fraction of intron 36 containing pre-mRNAs are not found at, but rather near, the site of transcription, indicating that splicing of this intron mostly occurs post-transcriptionally. Most of the nucleoplasmic intron 36 containing pre-mRNAs are also organized in a looped conformation, and this conformation is dependent on the assembly of U2 snRNP. This suggests that at least some spliceosomal components might already be assembled on these nucleoplasmic pre-mRNAs and that intron organization and spliceosome assembly could be coordinated co-transcriptionally for both co- and post-transcriptional splicing. Such as scenario might also suggest the existence of regulatory steps for the catalytic steps of splicing or spliceosome assembly that would occur at a step past complex B formation. Whether these observations are universal and applicable for other introns, especially for introns that are spliced slowly or pre-mRNAs containing detained/retained introns, or introns within lncRNAs, are interesting questions for future studies.

### Mechanisms for splicing of long introns

Our data also allowed us to investigate mechanisms through which long introns are spliced. One prominent model proposed to facilitate the splicing of long introns is recursive splicing (RS), which suggests that long introns be removed in multiple steps, avoiding the need to bring the ends of introns, sometimes separated by hundreds of thousands of nucleotides, to assemble the spliceosome. While RS is relatively frequent and has been well characterized in Drosophila, its extent in splicing of long introns in humans remains unclear, although recent transcriptome-wide studies suggest that it occurs on various introns (Sibley, Blazquez, and Ule 2016; Wan et al. 2021). Our data do not suggest recursive splicing to occur for POLA1 introns 35 and 36, as single molecules lacking signals from the 5’ end, while containing signals from the 3’ end, an indication that these introns are spliced in multiple steps, are extremely rare. Additionally, the presence of fully transcribed long introns such as POLA1 intron 35 and 36 in a conformation capable of undergoing splicing and with possibly at least partially assembled spliceosomes suggests that recursive splicing alone is unlikely to explain the splicing of all long introns in humans.

Alternatively, facilitating splice-site proximity at long introns has been proposed to be mediated by the tethering of 5’ ss bound U1 snRNP to the elongating RNA Pol II (Hollander et al. 2016; Zhang et al. 2021). We observe that the organization of intron 35 is altered during transcription, with distances between 5’ and 2/3^rd^ regions shorter for partially transcribed introns lacking the 3’ signal compared to introns that have been fully transcribed. These observations are compatible with a tethering model and could provide the mechanism through which the ends of POLA1 intron 35 and 36 are brought together once the polymerase reaches the 3’ end of the intron, resulting in the observed looped intronic conformations, which then become stabilized by the assembly of the spliceosome. Otherwise, a compact nascent intronic RNP could also be assembled in a way that results in a rigid intronic RNP that positions and maintains the introns 5’ end in close proximity to the emerging nascent RNA and any potential emerging 3’ ss, independent of polymerase tethering. Further characterization of the intronic RNP composition and biophysical properties and effects in disrupting the recently described interactions between U1 snRNP and components of the transcription machinery will be necessary to further dissect the mechanisms that allow efficient splicing of long introns.

## Materials and Methods

### Reagents used

Splicing inhibitor Pladienolide B was bought from Cayman Chemicals(#16538) – stock at 100 µM in DMSO. The drug was diluted in warm media to get its final working concentration of 100 nM, and cells were treated for four hours before fixation.

### Cell culture

HEK293T (American Type Culture Collection CRL-3216) cells were maintained at 37°C and 5% CO_2_ in Dulbecco’s Modified Eagle Medium (DMEM) (Wisent, 319-015-CL) supplemented with 10% fetal bovine serum (FBS) (Wisent, 080-150) and passaged every 2-3 days with Trypsin (Wisent 325-043-EL). Prior to the day of treatment and fixation, cells were plated on poly-L-Lysine (Sigma, P8920 - final concentration of 0.01% w/v) coated coverslips. On the day of treatment, the media was replaced with media containing the drug and placed back in the incubator. After incubation, the cells were briefly rinsed with 1x PBS, after which they were fixed using 4% PFA, 1X PBS for 10 min at room temperature. After that, the cells were washed twice with 1X PBS for 5 min each and permeabilized with ice-cold 70% ethanol. The coverslips were stored at - 20 °C for at least 12 hrs before being used for smFISH.

### smRNA FISH probe design and labelling

The sequences for the introns and mRNAs were obtained from Ensembl, and the designed probes are listed in Table S 1, and their distribution is shown in Supplementary Figure 1. Probe combinations used for experiments and distance measurements are listed in Table S 2 and mentioned in the figure legends. smFISH was done as previously described in (Adivarahan and Zenklusen 2021). Custom DNA probe sets targeting different regions of RNAs of interest were designed using Stellaris^®^ Probe Designer and either synthesized by Biosearch Technologies containing 3’ amine-reactive group or ordered as DNA oligos from Biobasic and ThermoFisher and modified in house by the addition of amine-modified ddUTP as previously described in Gaspar et al. (Gaspar, Wippich, and Ephrussi 2017). The amine-modified probes were labelled with far-red dye Cy5 (GEPA25001), orange dyes Cy3 (GEPA23001) from Sigma or Dylight 550 (Thermo Scientific 62263) or green dyes Dylight488 (Thermo Scientific 46403) or Atto488 (Thermo Scientific A20000) as previously described in (Adivarahan and Zenklusen 2021).

### smFISH

Prior to hybridization, cells were rehydrated in 1xPBS, then rinsed with 10% formamide/2xSSC for 10 mins at room temperature. The cells were hybridized with 10-20 ng (1.2-2.5 pmol) of each probe mix along with 40 μg of ssDNA/tRNA. The probes were resuspended in the hybridization solution (10% dextran sulfate/10% formamide/2xSSC/2 mM VRC/0.1 mg/ml BSA) and incubated for 3 hrs in the dark at 37°C. Post hybridization, the coverslips were washed 2x with 10% formamide/2xSSC solution for 30 min at 37 °C. The second wash was carried out in the presence of 0.5µg/ml. Samples were then rinsed with 1xPBS and mounted using ProLong Gold antifade reagent with DAPI (P36935, Invitrogen).

### Image Acquisition and pixel shift correction

Images were acquired with a 63x NA 1.46 oil objective on a Zeiss Elyra PS.1 system equipped with an Andor EMCCD iXon3 DU-885 CSO VP461 camera (1004×1002 pixels), and the following filter sets: BP420-480 + LP750 (Zeiss SR cube 07), BP495-590+LP750 (Zeiss SR cube 13), LP570 (Zeiss SR cube 14), LP655 (Zeiss SR cube 10) and the following lasers: 50 mW 405 nm HR diode, 100 mW 488 nm HR diode, 100 mW 561 nm HR DPSS, 200 mW 639 nm HR diode. Each image was acquired using three rotations and a grid size of 42 µm for all channels. The channels were registered using coverslips containing 0.1 µm TetraSpec beads (Invitrogen T-7279). Images for the beads were acquired in all channels, and the correction between channels was calculated and corrected using the built-in channel alignment tool in ZEN 2012 SP5 using the affine transformation. This correction was calculated for each day of imaging.

### RNA spot detection, spot assignment and distance measurements

3D images were processed using ImageJ, and the spots in different channels were separated using custom ImageJ scripts. 3D spot detection was carried out using FISH-Quant (Mueller et al. 2013). Only the X and Y coordinates were used from here on for further analysis. Masks were created in FiJi by manual segmentation as described in Adivarahan and Zenklusen 2021 for either separating nuclear and cytoplasmic RNAs or assigning spots corresponding to a single RNA. Assignment of the 5’, 3’ and/or the mid spots to either the cytoplasmic or the nuclear masks was done using MATLAB (MathWorks). To measure distances between different regions of mRNPs, spots from different channels were first grouped to assign neighbouring spots corresponding to a single RNA. This was achieved by using spots from one channel as a reference and finding spots from the other channels within a defined radius using the coordinates from the Gaussian fitted spots as described in Adivarahan and Zenklusen 2021. Alternatively, spots from different channels corresponding to a single mRNA were assigned manually within the mask. 2D distances between different regions of the RNA were then calculated for signals within a group.

### Data Plotting

All measurements were made for at least two independent biological replicates and the data plotted are representative from one of the replicates. For each measurement, at least ten different fields were imaged, with each image containing a minimum of 15 cells. A minimum of 200 RNAs was analyzed for each dataset unless stated otherwise. For partially transcribed intron 35, a total of 61 introns were analyzed from 36 different fields, primarily due to the lack of abundance of this intron. The centre of mass plots in Supplementary Figure 5B and Supplementary Figure 6E were made in R. The centre of mass was calculated as the mean of the coordinates of the three regions. The different conformations were then aligned using their centre of masses. The mean radius of gyration (<R_g_>) was calculated using:

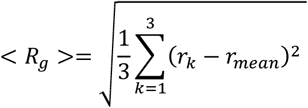

where k represents one of the three regions of the mRNP and r_k_ the position of the corresponding position in space as determined by 3D Gaussian fitting, but using the X and Y coordinates. The angle formed between the three regions was calculated using the 2D coordinates from 3D Gaussian fitting of individual introns. All plots were made in R.

## Acknowledgement

We thank members of the Zenklusen laboratory for critical discussion and comments on the project and the Nicolas Stifani at the BMM microscopy platform for support with the Zeiss ELYRA microscope. This work has been supported CIHR (Project Grant-366682), FRQ-S (Chercheur-boursier senior -DZ), FRQ-S (Formation de doctorat-SA) and CFI (DZ)

## Author contributions

SA and DZ conceived the study; SA developed the image analysis toolbox, performed and analyzed the smFISH experiments with the assistance of AMSKA. DZ supervised the work. SA and DZ wrote the manuscript.

