## Supplementary Figures and Tables for "Single-molecule imaging suggests compact and spliceosome dependent organization of long introns"

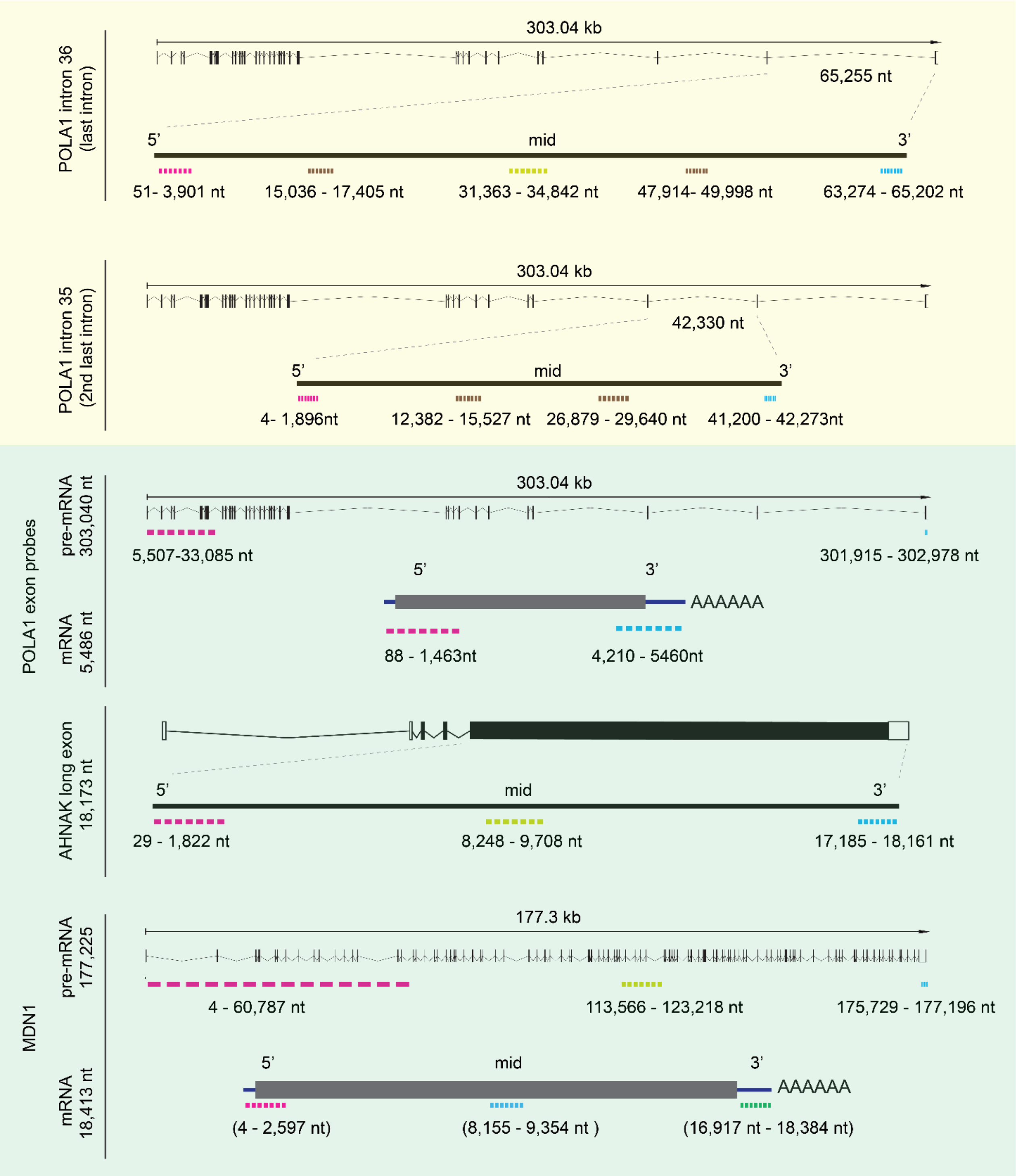


**Supplementary Figure 1: Positions of smFISH probes used in this study**

*Cartoons illustrating the positions of the probes used for the different genes used. See Table S 1 for probe sequences. The intron and transcript sequences were obtained from ensembl.*

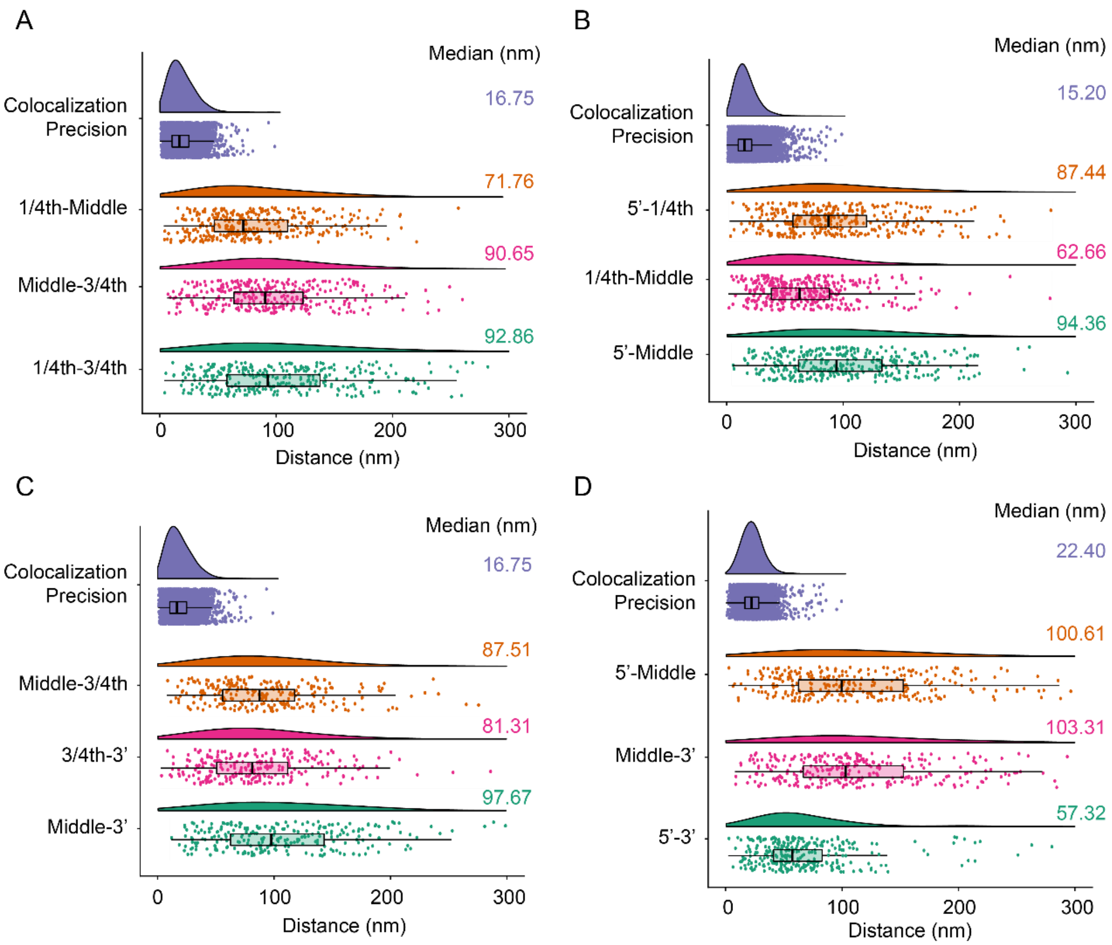


**Supplementary Figure 2: Intron 36 smFISH distance distributions**

*Raincloud plots for distances between different regions used in Figure 1 E. Individual plots represent individual smFISH experiments using the probe combinations as follows: Top-left: Probe Set#9, Table S 2, Top-right: Probe Set#8, Table S 2, Bottom-left: Probe Set#10, Table S 2 and Bottom-right: Probe Set#2, Table S 2. Individual plots show distance distribution of co-localization precision and distances for POLA1 introns as violin plots. The box plot shows the first quartile, median and third quartile and the distances corresponding to single RNAs are shown as spots overlayed on top of the box plots. Median distances are shown on the right*

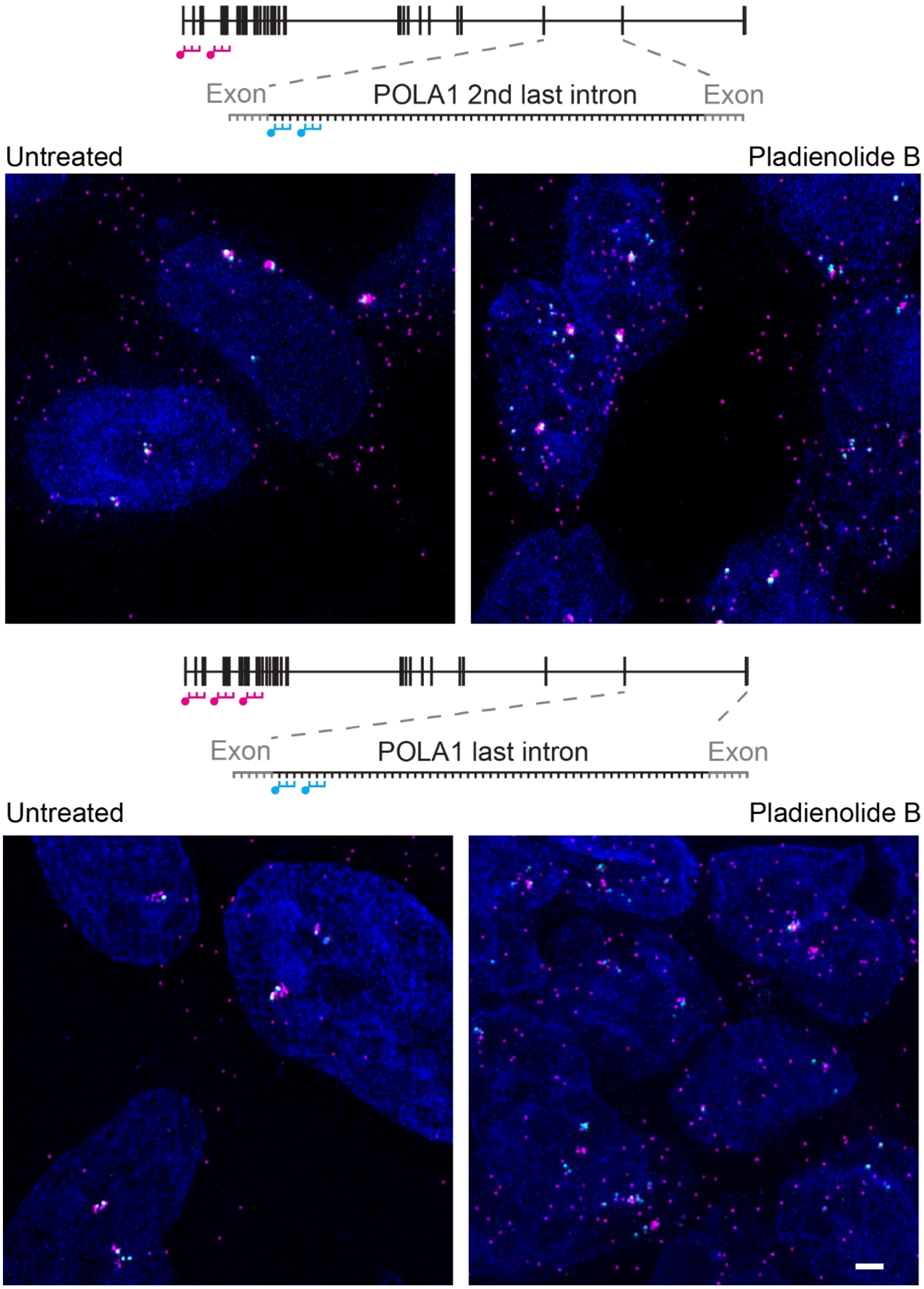


**Supplementary Figure 3: Splicing inhibition upon Pladienolide B treatment**

*smFISH images in mock HEK293T cells and cells treated with 100 nM Pladienolide B for 4h with probes hybridizing to the 5' exon and 5' end of the intron 35 (Probe Set#4, Table S 2) (top) or 5' end of intron 36 (bottom) (Probe Set#16, Table S 2). Nuclei are visualized by DAPI staining (blue). Schematic position of probes shown on top. Scale bar is 2 µm*

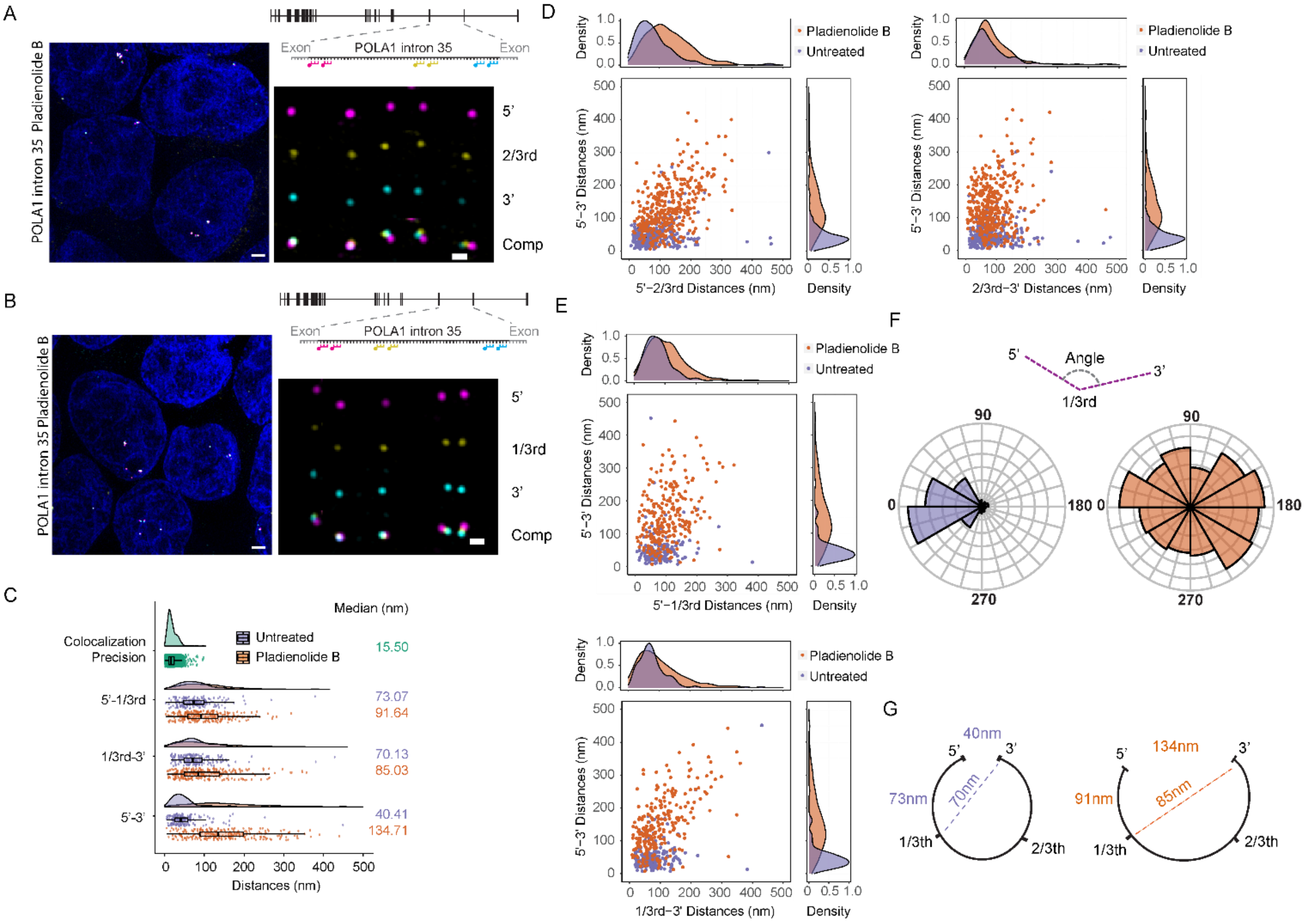


### Supplementary Figure 4: Pladienolide B treatment alters intron organization

***(A-B)*** *smFISH images in HEK293T cells treated with 100 nM Pladienolide B for 4h. Nuclei are visualized by DAPI staining (blue). Magnified images of individual RNAs are shown on the right. Schematic position of probes shown on top. Probes hybridizing to 5'(magenta), 3' (cyan) regions and either 2/3$^{rd}$(A) or 1/3$^{rd}$(B) regions (Probes Set#14 and #13 respectively, Table S 2).* ***(C)*** *Raincloud plots for distances between different regions for untreated and treated cells. Individual plots show distance distribution of co-localization precision and distances for POLA1 intron 35 as violin plots. The box plot shows the first quartile, median and third quartile and the individual RNAs shown as spots. Median distances are shown on the right.* ***(D-E)*** *Scatter plot showing distances for individual introns for intron 35 for mock and Pladienolide B treated HEK293T cells.* ***(F)*** *Angle histogram plot showing angle formed between the three regions of intron 35 for mock and Pladienolide B treatment as represented in the cartoon.* ***(G)*** *Cartoon representation of median* *distances between different regions of intron 35 as observed in C. Scale bars, 2 µm in larger images, and 500 nm in zoomed-in images*

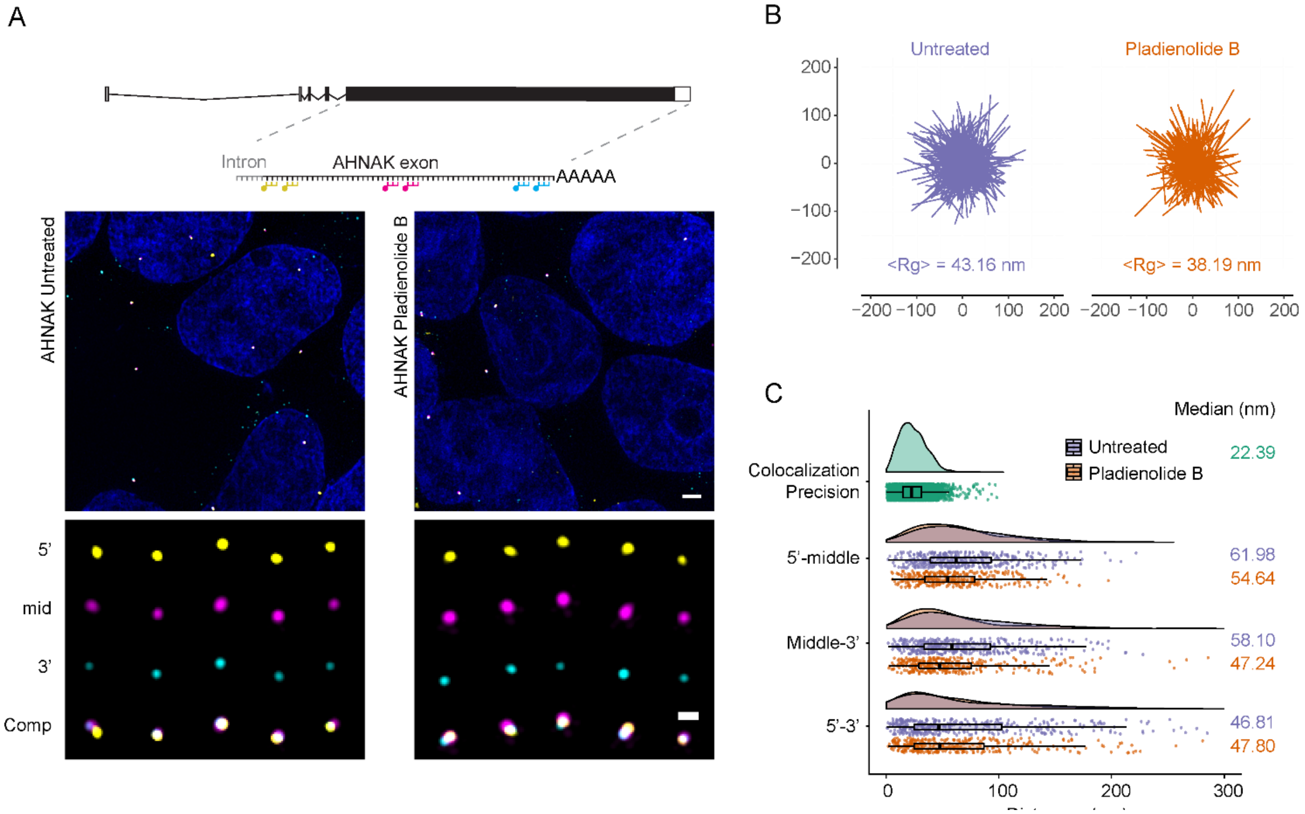


**Supplementary Figure 5: Organization of long AHNAK exon**

***(A)*** *smFISH images in HEK293T cells for probes hybridizing to 5'(yellow), middle (magenta), 3' (cyan) regions of AHNAK mRNAs (Probes Set#15, Table S 2).in mock and treated with 100nM Pladienolide B for 4h. Nuclei are visualized by DAPI staining (blue). Magnified images of individual RNAs are shown at the bottom. Schematic position of probes shown on top,* ***(B)*** *Projections of superimposed conformations from A with their centres of mass. The radius of gyration is shown at the bottom* ***(C)*** *Raincloud plots for distances between different regions used in A. Individual plots show distance distribution of colocalization precision and distances for AHNAK in mock and Pladienolide B treated cells shown as violin plots. The box plot shows the first quartile, median and third quartile and the individual RNAs shown as spots overlayed on top of the box plots. Median distances are shown on the right. Scale bars, 2 µm in larger images, and 500 nm in zoomed-in images*

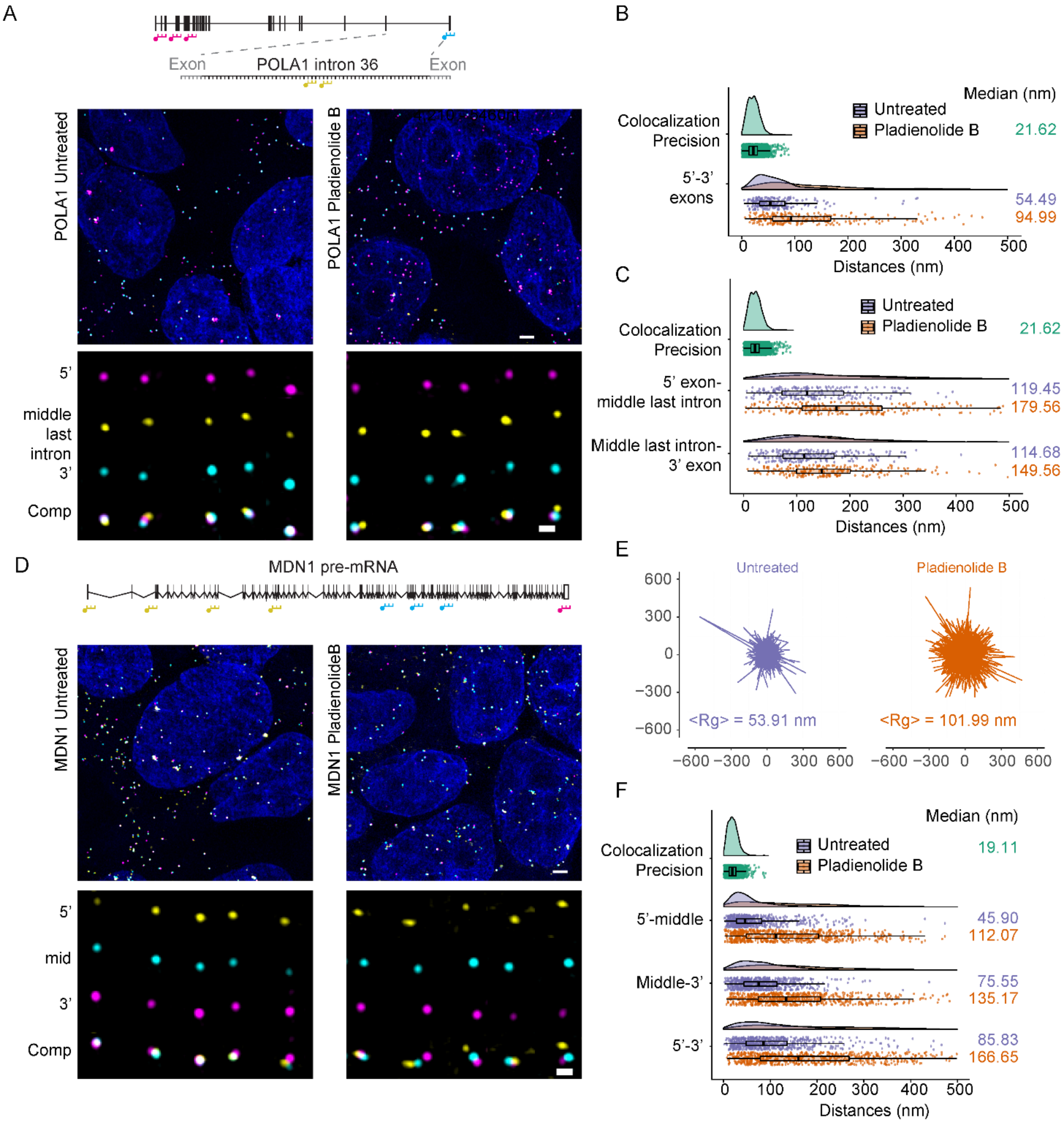


**Supplementary Figure 6: mRNA organization upon Pladienolide B treatment**

***(A, D)*** *smFISH images in HEK293T cells in mock and treated with 100nM Pladienolide B for4h. Nuclei are visualized by DAPI staining (blue). Magnified images of individual RNAs are shown at the bottom. Schematic position of probes shown on top, Probes hybridizing to* ***(A)*** *5’ exons (magenta), 3’exons (cyan) and middle region of intron 36 (yellow) of POLA1 gene (Probes Set#5,*

*Table S 2). and (D) to 5'(yellow), middle (magenta), 3' (cyan) exonic regions of MDN1 mRNAs (Probes Set#2, Table S 2).* ***(B,C,F) )*** *Raincloud plots for distances between different regions used in (A),(D). Individual plots show distance distribution of co-localization precision and distances between different regions as violin plots. The box plot shows the first quartile, median and third quartile and the individual RNAs shown as spots overlayed on top of the box plots. Median distances are shown on the right.* ***(E)*** *Projections of superimposed conformations from D with their centres of mass. The radius of gyration is shown below. Scale bars, 2 µm in larger images, and 500 nm in zoomed-in images*

### Supplementary Tables

#### Table S 1: List of smFISH probes used

| | | | |
|---|---|---|---|
| MDN1 exons 5’ | tcgttcttggctgcgattaa | POLA1 intron 36 tiling | gtgaaaaagcagttagtggc |
| MDN1 exons 5’ | taaggtactcaggacacact | POLA1 intron 36 tiling | agggtgacctgaagaatagt |
| MDN1 exons 5’ | cacagtacagtccttatcca | POLA1 intron 36 tiling | cccatagcactttgtgagaa |
| MDN1 exons 5’ | agcaaatccaaaaggagagg | POLA1 intron 36 tiling | cagcaacagtttttcgatgg |
| MDN1 exons 5’ | ttgaaagactggggatgtgt | POLA1 intron 36 tiling | gtgtgctttacctgttatta |
| MDN1 exons 5’ | gcatctgaactctctaggaa | POLA1 intron 36 tiling | aaacttgggtagtccttctg |
| MDN1 exons 5’ | gtgctcttcattcatacagg | POLA1 intron 36 tiling | gccaggttgtgatatgaatt |
| MDN1 exons 5’ | cctcaacctgaaatggatca | POLA1 intron 36 tiling | ctttctcgttcttgtagtga |
| MDN1 exons 5’ | aaaaccaaggccttctccaa | POLA1 intron 36 tiling | atgctgccatcagacaatac |
| MDN1 exons 5’ | aaagggagacttctggattg | POLA1 intron 36 tiling | ccactggaatagccttttaa |
| MDN1 exons 5’ | tcagacgaaacaagatgtcc | POLA1 intron 36 tiling | aaccatggaaagccttccac |
| MDN1 exons 5’ | accagcacataagacctaag | POLA1 intron 36 tiling | agtatacctgtatctaaccc |
| MDN1 exons 5’ | gaagacttttgcagacagac | POLA1 intron 36 tiling | agcactaattttggggagga |
| MDN1 exons 5’ | acagcattctgagaagcaac | POLA1 intron 36 tiling | tgatgggctactaagtctgt |
| MDN1 exons 5’ | tcctattggtccttccaaca | POLA1 intron 36 tiling | ggcctttatacagggagaat |
| MDN1 exons 5’ | cctgtcactgcagctaaata | POLA1 intron 36 tiling | atttctttcaaagcaccacc |
| MDN1 exons 5’ | aagctggactttgagaagct | POLA1 intron 36 tiling | gatgtatcctgtaggatgga |
| MDN1 exons 5’ | acatctgtgcagcgatacat | POLA1 intron 36 tiling | acatggtagggtaattcctg |
| MDN1 exons 5’ | atatcctccagaaggatcca | POLA1 intron 36 tiling | tagtccatgtctggaatagc |
| MDN1 exons 5’ | aagagctctccattctccaa | POLA1 intron 36 tiling | cattttgaggacactgcttc |
| MDN1 exons 5’ | aaatccaggtgccactttca | POLA1 intron 36 tiling | gcttacatcatttgagccaa |
| MDN1 exons 5’ | caactacgcccgcaggaaag | POLA1 intron 36 tiling | atgcttcctttatctaggaa |
| MDN1 exons 5’ | agcaagaagtgctccatgac | POLA1 intron 36 tiling | gctacattagcgtatacagc |
| MDN1 exons 5’ | caagaacctgcccaactcac | POLA1 intron 36 tiling | gaaaccacaccagccaaagg |
| MDN1 exons 5’ | cgatcttgaggtgtccacac | POLA1 intron 36 tiling | tgggttgaggttggctaacc |
| MDN1 exons 5’ | aatggcttcggcattccttt | POLA1 intron 36 tiling | atctggagcagagagctaaa |
| MDN1 exons 5’ | gttcatgcagatcatggttg | POLA1 intron 36 tiling | aggctagccaaccaaataac |
| MDN1 exons 5’ | gtttgctcatcgacacacat | POLA1 intron 36 tiling | cagcctttttattcagttga |
| MDN1 exons 5’ | gacatcaggatggttaccaa | POLA1 intron 36 tiling | cacccaaaggaatcagcagt |
| MDN1 exons 5’ | aatatctcagggcaaacggg | POLA1 intron 36 tiling | acatgcctcaagatattttt |
| MDN1 exons 5’ | tacgtccatagcgtactgga | POLA1 intron 36 tiling | ggagggggctggttaaaaaa |
| MDN1 exons 5’ | gcagaaacttgaaggctgct | POLA1 intron 36 tiling | tggaaggcaatctaagcaga |
| MDN1 exons 5’ | cggaacacagactgctcctg | POLA1 intron 36 tiling | gctacctgtgtatgattcaa |
| MDN1 exons 5’ | aaggtgtcatggcttctgag | POLA1 intron 36 tiling | aatgtactggtgaagtgggg |
| MDN1 exons 5’ | acaattggctgtataccagc | POLA1 intron 36 tiling | ataaacctgtcactgcagtg |
| MDN1 exons 5’ | acaccacaaacagctgtcac | POLA1 intron 36 tiling | gacatcacatagcaaggctt |
| MDN1 exons 5’ | tcttactgtcagtctgatct | POLA1 intron 36 tiling | tagctgagccatgtcataac |
| MDN1 exons 5’ | gaagaactcctattaccacc | POLA1 intron 36 tiling | aatacacagagtgcttccag |

| | | | |
|---|---|---|---|
| MDN1 exons 5’ | ctgccacacaaactctccag | POLA1 intron 36 tiling | gacctccgtacataatttct |
| MDN1 exons 5’ | cagaaaccacgtctaagggg | POLA1 intron 36 tiling | ggtacagtcacaatcatgca |
| MDN1 exons 5’ | caatttcctccacagctcaa | POLA1 intron 36 tiling | tatgcacccattacagatga |
| MDN1 exons 5’ | atgactgtttagcggtcgat | POLA1 intron 36 tiling | aattgtgttcctgagtcact |
| MDN1 exons 5’ | ccaggtgaattttggtccaa | POLA1 intron 36 tiling | acaatggggactgcctaatg |
| MDN1 exons 5’ | cagttctctcttatccaggt | POLA1 intron 36 tiling | taaactatgccctgcatcta |
| MDN1 exons 5’ | gtttctctccagtaagttgg | POLA1 intron 36 tiling | gccaaaattctattctagcg |
| MDN1 exons 5’ | gaactatcactccaagagtg | POLA1 intron 36 tiling | tatacagctgaggctagagg |
| MDN1 exons 5’ | aacttcttcaggtgcctgtt | POLA1 intron 36 tiling | atttttactcttaacctccc |
| MDN1 exons 5’ | ctcaagggttggtcttttgt | POLA1 intron 36 tiling | gccagtctttttctaaactg |
| MDN1 exons 5’ | gggcaatcctattacaccaa | POLA1 intron 36 tiling | aaatgactgctggcatctca |
| MDN1 exons 5’ | ctcagaaagcattgctgtga | POLA1 intron 36 tiling | ctttatcccaaaagacttca |
| MDN1 exons 5’ | gcttccaataacttctgcca | POLA1 intron 36 tiling | gaaggcatggtgtgttaagc |
| MDN1 exons 5’ | tttgctgacacacactgcaa | POLA1 intron 36 tiling | ctagtgaatgtcctcactga |
| MDN1 exons 5’ | atggtagaggttttgccagt | POLA1 intron 36 tiling | attgtgctggttaggataca |
| MDN1 exons 5’ | cctgtaatgtgagccaagta | POLA1 intron 36 tiling | agagacttgccattaagtgg |
| MDN1 exons 5’ | attgacaaccctcaaacggt | POLA1 intron 36 tiling | acagcagttgactgttgaga |
| MDN1 exons 5’ | gcaagtctgcagtatcactt | POLA1 intron 36 tiling | attcaacctgtcaaagctgc |
| MDN1 exons 5’ | atggtccaccggtttataac | POLA1 intron 36 tiling | gaacgtatcacccactattg |
| MDN1 exons 5’ | gtaagggtagccaaataagc | POLA1 intron 36 tiling | atgatggttaatgggaaggt |
| MDN1 exons 5’ | aagagttcctcaaatgcctc | POLA1 intron 36 tiling | ctgggtattatagttacctc |
| MDN1 exons 5’ | gccccaagaacgtaaagttt | POLA1 intron 36 tiling | ttctcaccttgcttttttca |
| MDN1 exons 5’ | ctgtctgtaacaggtctgaa | POLA1 intron 36 tiling | tttcttatccctgacttcag |
| MDN1 exons 5’ | ttagtctcaggagatcatgc | POLA1 intron 36 tiling | cgtttcttgtgtttttgacc |
| MDN1 exons 5’ | agcagacttgtgtacatgct | POLA1 intron 36 tiling | ctgcgtgtgaagctcaaatt |
| MDN1 exons 5’ | cactgtcttttccatccttg | POLA1 intron 36 tiling | agccacactattatatgttt |
| MDN1 exons 5’ | ccaaatgcttcccatttctc | POLA1 intron 36 tiling | aattcacccttttacactctc |
| MDN1 exons 5’ | ttgggcatggttgagtctaa | POLA1 intron 36 tiling | gatctgccaacataccttac |
| MDN1 exons middle | aagcagcaagattgaccaca | POLA1 intron 36 tiling | aacatgatcacggactccac |
| MDN1 exons middle | ggactagtgcatcacaaagt | POLA1 intron 36 tiling | aaatactaggtctcctctct |
| MDN1 exons middle | actggcatcagacaccattc | POLA1 intron 36 tiling | aaaggctcatttttcccatg |
| MDN1 exons middle | accgcagagaacctaagatc | POLA1 intron 36 tiling | gtgtctagaaaaccattgct |
| MDN1 exons middle | ggcatctacttttactgtgt | POLA1 intron 36 tiling | caaagaccaaggccattttc |
| MDN1 exons middle | tgccaatggagggcaagaag | POLA1 intron 36 tiling | ggtagctgaatttgctgaga |
| MDN1 exons middle | ctggtggaccagatgtttta | POLA1 intron 36 tiling | accttagctctagaactctt |
| MDN1 exons middle | aattcatcagaagtcggggg | POLA1 intron 36 tiling | aaggcatgctgtgccaagac |
| MDN1 exons middle | ctgagacagtctgaacttct | POLA1 intron 36 tiling | ggcttattctcttgatgagg |
| MDN1 exons middle | tccccagacaattctgaata | POLA1 intron 36 tiling | ccataaatcattagccttgt |
| MDN1 exons middle | cttctttataccagcgaagc | POLA1 intron 36 tiling | ttggggacatcatctttagt |
| MDN1 exons middle | aaacggtcgtcccaggaact | POLA1 intron 36 tiling | aaatgcatagtcgtcacctg |
| MDN1 exons middle | ccaccagcttgtccttaaaa | POLA1 intron 36 tiling | ggtaagagtcggatctgtta |
| MDN1 exons middle | ggaccttcagttgtgaaaaa | POLA1 intron 36 tiling | aaacctgacagaggtgggtg |

| | | | |
|---|---|---|---|
| MDN1 exons middle | ctgatggcaaggactttgtt | POLA1 intron 36 tiling | accaataacgtagcatagct |
| MDN1 exons middle | agacggttaatgtcttcctg | POLA1 intron 36 tiling | ggcataatgcccactttaaa |
| MDN1 exons middle | ccactgagaagcaaccactt | POLA1 intron 36 tiling | agattggttttccattgtgc |
| MDN1 exons middle | cttgcaggagacttttcttt | POLA1 intron 36 tiling | aaattgcgtttgggcttctc |
| MDN1 exons middle | atttgctctgaggatcagtc | POLA1 intron 36 tiling | accatatctttcactggcaa |
| MDN1 exons middle | ttcatctaggctgacatctt | POLA1 intron 36 tiling | aaactgaaactcgccatcct |
| MDN1 exons middle | ctgagcatgcacaaaattct | POLA1 intron 36 tiling | tgcacatctcagattcttct |
| MDN1 exons middle | cctttggctttcagttctaa | POLA1 intron 36 tiling | ccaaatggggggttattaga |
| MDN1 exons middle | ctccagaaaaccaagtgaga | POLA1 intron 36 tiling | tactgactccaatggctgac |
| MDN1 exons middle | aggaagcttcatcatgcttt | POLA1 intron 36 tiling | cctcttttcaactcttctta |
| MDN1 exons middle | gtcaagtctggatgggacag | POLA1 intron 36 tiling | ggttgatcactaacatagcc |
| MDN1 exons middle | tcctggtgaggtggattacg | POLA1 intron 36 tiling | caaggcactgtcaaacaggt |
| MDN1 exons middle | attgcaggccacaactgaac | POLA1 intron 36 tiling | atctgtggataacctgctac |
| MDN1 exons middle | tttgtaccgccaaagcatag | POLA1 intron 36 tiling | cagtttctgagaccatgtta |
| MDN1 exons middle | gtgccataaaatcagctgtc | POLA1 intron 36 tiling | cacaggctgacggattagag |
| MDN1 exons middle | ctgcatcttctgagacaagc | POLA1 intron 36 tiling | acatattcaaccagtgttca |
| MDN1 exons middle | tttatctggggttgttgatt | POLA1 intron 36 tiling | atcctgggaagtttatttgt |
| MDN1 exons middle | agatgaggtgacttatctcc | POLA1 intron 36 tiling | atttaactctgttacctgca |
| MDN1 exons middle | acaggtgtgtgataaagaca | POLA1 intron 36 tiling | ctgtgtttgaaattgctggt |
| MDN1 exons middle | atctcgaagctcttgaggtg | POLA1 intron 36 tiling | aatgctggaagacaagctta |
| MDN1 exons middle | tgatgatgcagcaaggacca | POLA1 intron 36 tiling | aacgaaagaggggactcggg |
| MDN1 exons middle | aatttcttcaggagacacct | POLA1 intron 36 tiling | ttagccaatgcctagaggac |
| MDN1 exons middle | ataactcggaccacaaagat | POLA1 intron 36 tiling | cagtgatcaatgtgtctatc |
| MDN1 exons middle | agtactgctccagaaagaca | POLA1 intron 36 tiling | agggtagcctagatactaat |
| MDN1 exons middle | agtactctggatttgtggtc | POLA1 intron 36 tiling | cccctatgaaaacgatggaa |
| MDN1 exons middle | ggcaaagggttccacattag | POLA1 intron 36 tiling | ctgtggaacttaatggccat |
| MDN1 exons middle | tccaaaacagacttgggtgc | POLA1 intron 36 tiling | agcaggtggtagcattatag |
| MDN1 exons middle | tgggtctattgagattgccg | POLA1 intron 36 tiling | acccacagttcaaagaacta |
| MDN1 exons middle | tcaaagcagcacttagagaa | POLA1 intron 36 tiling | cacagattctactattcctt |
| MDN1 exons middle | tctccagctgctggttaaga | POLA1 intron 36 tiling | gtgcctgcattagatcataa |
| MDN1 exons 3’ | aagtctcactttggactctt | POLA1 intron 36 tiling | tgacaccctgacatctgatc |
| MDN1 exons 3’ | aatgtgaccttctgaccaca | POLA1 intron 36 tiling | aattgctgcagtccattcac |
| MDN1 exons 3’ | aaaagggagcacctgggtaa | POLA1 intron 36 tiling | acaaggttaactaggttcgt |
| MDN1 exons 3’ | agcattctgtaggctgtaag | POLA1 intron 35 1/3rd | tcagtgtttgtgattatact |
| MDN1 exons 3’ | tggtataaaaacctcagccc | POLA1 intron 35 1/3rd | ttcacagtgtcactttcaca |
| MDN1 exons 3’ | actcttctctagttacgagt | POLA1 intron 35 1/3rd | atccccaaatgccttgaaaa |
| MDN1 exons 3’ | cttccaaggcagggaagaag | POLA1 intron 35 1/3rd | gtcctgttgaatgctgaact |
| MDN1 exons 3’ | aagaaaacaggcagctggg c | POLA1 intron 35 1/3rd | aaatgcatctgcctacatct |
| MDN1 exons 3’ | acaaaggactgtcagagtcc | POLA1 intron 35 1/3rd | aaggatagcacagagggcct |
| MDN1 exons 3’ | aaaagggcagctccctttag | POLA1 intron 35 1/3rd | ggaaggtaaatgcaggaaca |
| MDN1 exons 3’ | gcaaggcagagcttagaaca | POLA1 intron 35 1/3rd | ttgcactccattgcatagaa |
| MDN1 exons 3’ | tttgggcacacactatgggc | POLA1 intron 35 1/3rd | ttcaggactgccaataaaca |

| | | | |
|---|---|---|---|
| MDN1 exons 3’ | ctgtcttggccacttgacag | POLA1 intron 35 1/3rd | ccattcagtacctctgcaaa |
| MDN1 exons 3’ | cctcatactctccagaaacg | POLA1 intron 35 1/3rd | ggaggatttggaaatctcag |
| MDN1 exons 3’ | tctaagagaaggtagttcct | POLA1 intron 35 1/3rd | aaagagtgctgccctaaaac |
| MDN1 exons 3’ | ctataatgtccagttgcttt | POLA1 intron 35 1/3rd | atttgggactcagaaatcct |
| MDN1 exons 3’ | tttttatagatgacctgggc | POLA1 intron 35 1/3rd | tcaaggttatgtctgaagga |
| MDN1 exons 3’ | tttttacacagcccaaggat | POLA1 intron 35 1/3rd | cattccaaggacacacagcc |
| MDN1 exons 3’ | gaggatactgaaaagccact | POLA1 intron 35 1/3rd | agcataaaatgatctctgct |
| MDN1 exons 3’ | cattgcatagtctcccgaag | POLA1 intron 35 1/3rd | cgtcacaggggaagaaggat |
| MDN1 exons 3’ | ataaaggggcaatcaccttc | POLA1 intron 35 1/3rd | aaggaatctctcctccattt |
| MDN1 exons 3’ | tacaacaacagggaccatgg | POLA1 intron 35 1/3rd | attgagaatcagtagcctgt |
| MDN1 exons 3’ | agtgtgaggaatcactcttc | POLA1 intron 35 1/3rd | tccataggcaagatcttagt |
| MDN1 exons 3’ | tggctcagtccagcttgaaa | POLA1 intron 35 1/3rd | agtaaggatactgctatgct |
| MDN1 exons 3’ | cgtttctttcccagaatgag | POLA1 intron 35 1/3rd | atttttttcccctaaagtct |
| MDN1 exons 3’ | atagatggagctgctgagtt | POLA1 intron 35 1/3rd | ggctagactgtgagctgaaa |
| MDN1 exons 3’ | atcagtttctttcgactgga | POLA1 intron 35 1/3rd | ggtttctctggttctgtaat |
| MDN1 exons 3’ | ggccaagtaaaaactgccta | POLA1 intron 35 1/3rd | ctgtacatgcatgtacaggg |
| MDN1 exons 3’ | caagtattcagcactgcttt | POLA1 intron 35 1/3rd | gtgcttctcatgactttcag |
| MDN1 exons 3’ | agtagaacagagcacacagt | POLA1 intron 35 1/3rd | cacacccatctgagacaatg |
| MDN1 exons 3’ | atcatgacatactgcctaca | POLA1 intron 35 1/3rd | cttcaacacacctgcatatg |
| MDN1 exons 3’ | tcgcagacttcacagtgtaa | POLA1 intron 35 1/3rd | ttgttaattacttcccacca |
| MDN1 exons 3’ | atctgtgtctttgatgacca | POLA1 intron 35 1/3rd | gcagaagcacagagaaggg a |
| MDN1 exons 3’ | tacatgctttgggacacttg | POLA1 intron 35 1/3rd | gagtagaaggcaccaaatcc |
| MDN1 exons 3’ | aagatcagtcccctagcata | POLA1 intron 35 1/3rd | actggagcatcttctaaata |
| MDN1 exons 3’ | ctgactgactgatccagcag | POLA1 intron 35 1/3rd | cctcctaaggctgttttcat |
| MDN1 exons 3’ | gcacagcatcaactagtaac | POLA1 intron 35 1/3rd | ctgtttctaagtagatgtca |
| MDN1 exons 3’ | gaagtaggaggggatcatgt | POLA1 intron 35 1/3rd | gctctaaagatccaacttca |
| MDN1 exons 3’ | cctttgtagtaagagcaaca | POLA1 intron 35 1/3rd | actggagtctaacagtcaca |
| MDN1 exons 3’ | cagcctaccatggacataaa | POLA1 intron 35 1/3rd | gaagggcatagtctgatacc |
| MDN1 exons 3’ | ctgcaaagccagcatattat | POLA1 intron 35 1/3rd | cctttaatctataaccatcc |
| MDN1 exons 3’ | gcctccttataaggctacac | POLA1 intron 35 1/3rd | atttcaagatttgccctaga |
| MDN1 middle Odd | aagcagcaagattgaccaca | POLA1 intron 35 1/3rd | aacagagtatggagcacacc |
| MDN1 middle Odd | actggcatcagacaccattc | POLA1 intron 35 1/3rd | ggaaggacaaggcataagga |
| MDN1 middle Odd | ggcatctacttttactgtgt | POLA1 intron 35 1/3rd | ggaagaggagagggaagac t |
| MDN1 middle Odd | ctggtggaccagatgtttta | POLA1 intron 35 1/3rd | ggtgctggttaatcatcaag |
| MDN1 middle Odd | ctgagacagtctgaacttct | POLA1 intron 35 1/3rd | caataaagcagggagacagt |
| MDN1 middle Odd | cttctttataccagcgaagc | POLA1 intron 35 1/3rd | tacgagtgtctttgtacatt |
| MDN1 middle Odd | ccaccagcttgtccttaaaa | POLA1 intron 35 1/3rd | atgttccaatcagtctttct |
| MDN1 middle Odd | ctgatggcaaggactttgtt | POLA1 intron 35 1/3rd | caggatgctggagatagaca |
| MDN1 middle Odd | ccactgagaagcaaccactt | POLA1 intron 35 1/3rd | aacgttttttgtttcccacg |
| MDN1 middle Odd | atttgctctgaggatcagtc | POLA1 intron 35 1/3rd | aaaggcagaacccattgttt |
| MDN1 middle Odd | ctgagcatgcacaaaattct | POLA1 intron 35 1/3rd | cttagttccccatattcaag |

| | | | |
|---|---|---|---|
| MDN1 middle Odd | ctccagaaaaccaagtgaga | POLA1 intron 35 $1/3^{rd}$ | gttgttttgcttcaaaggtc |
| MDN1 middle Odd | gtcaagtctggatgggacag | POLA1 intron 35 $1/3^{rd}$ | ctcttcagcactgatgtatt |
| MDN1 middle Odd | attgcaggccacaactgaac | POLA1 intron 35 $1/3^{rd}$ | ctttactaactactttaccc |
| MDN1 middle Odd | gtgccataaaatcagctgtc | POLA1 intron 35 $1/3^{rd}$ | ccttactaactactttatcc |
| MDN1 middle Odd | tttatctggggttgttgatt | POLA1 intron 35 $2/3^{rd}$ | ccagtaattttaaaccagga |
| MDN1 middle Odd | acaggtgtgtgataaagaca | POLA1 intron 35 $2/3^{rd}$ | caccttgctcagatatactg |
| MDN1 middle Odd | tgatgatgcagcaaggacca | POLA1 intron 35 $2/3^{rd}$ | ccaatatgactcctctgtaa |
| MDN1 middle Odd | ataactcggaccacaaagat | POLA1 intron 35 $2/3^{rd}$ | atctgcaggcctgtaaggag |
| MDN1 middle Odd | agtactctggatttgtggtc | POLA1 intron 35 $2/3^{rd}$ | aagccactggaggcagtatg |
| MDN1 middle Odd | tccaaaacagacttgggtgc | POLA1 intron 35 $2/3^{rd}$ | aataggtgggtggggcttaa |
| MDN1 middle Odd | tcaaagcagcacttagagaa | POLA1 intron 35 $2/3^{rd}$ | aatactgttccccttcagaa |
| MDN1 middle Even | ggactagtgcatcacaaagt | POLA1 intron 35 $2/3^{rd}$ | gtatgtgaatttgctaagcc |
| MDN1 middle Even | accgcagagaacctaagatc | POLA1 intron 35 $2/3^{rd}$ | agcaaattggtagatcatca |
| MDN1 middle Even | tgccaatggagggcaagaag | POLA1 intron 35 $2/3^{rd}$ | ttatgctcttcaacacaggc |
| MDN1 middle Even | aattcatcagaagtcggggg | POLA1 intron 35 $2/3^{rd}$ | ccacagaatgcagcttcaaa |
| MDN1 middle Even | tccccagacaattctgaata | POLA1 intron 35 $2/3^{rd}$ | ggatatctttggatagagcc |
| MDN1 middle Even | aaacggtcgtcccaggaact | POLA1 intron 35 $2/3^{rd}$ | gaaccccttttcttttagat |
| MDN1 middle Even | ggaccttcagttgtgaaaaa | POLA1 intron 35 $2/3^{rd}$ | gttcacaatattttcttggt |
| MDN1 middle Even | agacggttaatgtcttcctg | POLA1 intron 35 $2/3^{rd}$ | tttcatttgtccaggtgagt |
| MDN1 middle Even | cttgcaggagacttttcttt | POLA1 intron 35 $2/3^{rd}$ | ctcaagttctcattatctgt |
| MDN1 middle Even | ttcatctaggctgacatctt | POLA1 intron 35 $2/3^{rd}$ | ggaaatggggatgactggtt |
| MDN1 middle Even | cctttggctttcagttctaa | POLA1 intron 35 $2/3^{rd}$ | ttttgctgttagcaacactt |
| MDN1 middle Even | aggaagcttcatcatgcttt | POLA1 intron 35 $2/3^{rd}$ | agataaagtaatgcaccccc |
| MDN1 middle Even | tcctggtgaggtggattacg | POLA1 intron 35 $2/3^{rd}$ | ctcagcagtttttccaaagg |
| MDN1 middle Even | tttgtaccgccaaagcatag | POLA1 intron 35 $2/3^{rd}$ | gataatccatgtcctcattc |
| MDN1 middle Even | ctgcatcttctgagacaagc | POLA1 intron 35 $2/3^{rd}$ | gtggcactcaattactctga |
| MDN1 middle Even | agatgaggtgacttatctcc | POLA1 intron 35 $2/3^{rd}$ | ctggcacattagtcagtaca |
| MDN1 middle Even | atctcgaagctcttgaggtg | POLA1 intron 35 $2/3^{rd}$ | gcccatttatctcatttgaa |
| MDN1 middle Even | aatttcttcaggagacacct | POLA1 intron 35 $2/3^{rd}$ | gcaggtctgtttctctgaaa |
| MDN1 middle Even | agtactgctccagaaagaca | POLA1 intron 35 $2/3^{rd}$ | acagaagtgcctcaatctga |
| MDN1 middle Even | ggcaaagggttccacattag | POLA1 intron 35 $2/3^{rd}$ | caatcctcttggctctaatt |
| MDN1 middle Even | tgggtctattgagattgccg | POLA1 intron 35 $2/3^{rd}$ | gcaggctgtgttaacatgaa |
| MDN1 middle Even | tctccagctgctggttaaga | POLA1 intron 35 $2/3^{rd}$ | ccaatatcttatctgattcc |
| POLA1 exons 3’ | tgtacagggacttgtcagaa | POLA1 intron 35 $2/3^{rd}$ | aattcttcaattcccaagcc |
| POLA1 exons 3’ | taccggtaaaagcacagctg | POLA1 intron 35 $2/3^{rd}$ | aggttcaaggatcaacaaca |
| POLA1 exons 3’ | gtgcacactccgcatcaaaa | POLA1 intron 35 $2/3^{rd}$ | cctctatctgggatttgaac |
| POLA1 exons 3’ | tctcatgatcggtagtaagt | POLA1 intron 35 $2/3^{rd}$ | catatttagatgccattcca |
| POLA1 exons 3’ | ctgtagtcctgcagaacttt | POLA1 intron 35 $2/3^{rd}$ | taataggcttcattctagcc |
| POLA1 exons 3’ | ctctgctgtgttcttgagtt | POLA1 intron 35 $2/3^{rd}$ | tacatcccattgatgttctc |
| POLA1 exons 3’ | tagccacttcgggacaagaa | POLA1 intron 35 $2/3^{rd}$ | aagtatgggaagtcctttga |
| POLA1 exons 3’ | tttgctcagattcacttcgg | POLA1 intron 35 $2/3^{rd}$ | tatacatctccccagtctag |
| POLA1 exons 3’ | ttaggatttcacggcacaac | POLA1 intron 35 $2/3^{rd}$ | cactcaattcctgtaacttt |

| | | | |
|---|---|---|---|
| POLA1 exons 3’ | cttggttactcctgggattc | POLA1 intron 35 2/3rd | gtgttctctctcaactactt |
| POLA1 exons 3’ | ggaagctgggatttttcaac | POLA1 intron 35 2/3rd | gctaatggattttccttgga |
| POLA1 exons 3’ | caagggagaaacagatgctg | POLA1 intron 35 2/3rd | tgctttctacatgcgataga |
| POLA1 exons 3’ | acacaaacatgagacacagt | POLA1 intron 35 2/3rd | tcccactgaaagcttaccta |
| POLA1 exons 3’ | gactcaacatttttgcagcc | POLA1 intron 35 2/3rd | gcttgtatacaagatctggt |
| POLA1 exons 3’ | ctcagaaaccgggtcttcag | POLA1 intron 35 2/3rd | gcactgcaaaaacagagcca |
| POLA1 exons 3’ | gctactctcaatccaagtag | POLA1 intron 35 2/3rd | attcactaattccttgtcta |
| POLA1 exons 3’ | gcctggggtcacttacattc | POLA1 intron 35 2/3rd | gtgattttcctccatgatat |
| POLA1 exons 3’ | cataggctaaaggccctgag | POLA1 intron 35 2/3rd | tactgaatcacggtagaggt |
| POLA1 exons 3’ | ttcagtcaggctctgagaag | POLA1 intron 35 2/3rd | ggtttggactttctttagga |
| POLA1 exons 3’ | tgaaaaagcaaaacgtcagc | POLA1 intron 35 2/3rd | gagatcatttagtggagtcc |
| POLA1 exons 3’ | ttagaccgggtttaattggc | POLA1 intron 35 2/3rd | ccagccatcataatgagaga |
| POLA1 exons 3’ | actcctggatggctggagaa | POLA1 intron 35 2/3rd | aggacacagactctctatcc |
| POLA1 exons 3’ | agacaagactgaaaaggaca | POLA1 intron 35 2/3rd | ctaaacatggctgctaggct |
| POLA1 exons 3’ | agtgcaaggcttctaaatct | POLA1 intron 35 2/3rd | ccagctcagtcaatgaaata |
| POLA1 exons 3’ | gagcaattcaacaaacaagc | POLA1 intron 36 1/4th | tgcagattattcgtgtttca |
| POLA1 exons 3’ | cagtgtgtgtctgttggact | POLA1 intron 36 1/4th | ctccctttctatatcaaagt |
| POLA1 exons 3’ | atgtgagtgtaaaacacctg | POLA1 intron 36 1/4th | attgttatctagatgccacc |
| POLA1 exons 3’ | gcactttctatttaaggggc | POLA1 intron 36 1/4th | gtgtttgacttcattttcct |
| POLA1 exons 3’ | ccctacacatgttaatggat | POLA1 intron 36 1/4th | gcttcctttatctaggaaat |
| POLA1 exons 3’ | tgaatacaacacagtgatcc | POLA1 intron 36 1/4th | ttagcgtatacagcaccata |
| POLA1 exons 3’ | ggccaattaagcatccctct | POLA1 intron 36 1/4th | ttcactttatctttgatgct |
| POLA1 exons 3’ | agaaaaaaatagcaagcgcc | POLA1 intron 36 1/4th | tacttttccctagcctattt |
| POLA1 exons 3’ | agacggtcctattgtgaaga | POLA1 intron 36 1/4th | ccagaaaacagttccttctt |
| POLA1 exons 3’ | aaatacattttgctgctgcc | POLA1 intron 36 1/4th | tttcatcttcagtttgctca |
| POLA1 exons 3’ | agagaggaaagactgccata | POLA1 intron 36 1/4th | gccaattttgttgaaatgca |
| POLA1 exons 3’ | cagcataattgtacaagggg | POLA1 intron 36 1/4th | gggggaaaataatgtgcttt |
| POLA1 exons 3’ | agaagaaggcacaacatact | POLA1 intron 36 1/4th | tattccttcattcatgacgc |
| POLA1 exons 5’ | aactccctgaatctgacaga | POLA1 intron 36 1/4th | tccagtctccttgaaagatc |
| POLA1 exons 5’ | ttgattttttttctcgccgg | POLA1 intron 36 1/4th | cttaatatctccccagttac |
| POLA1 exons 5’ | tttctagggcttcttggcgc | POLA1 intron 36 1/4th | aagtaccacgtttgggttag |
| POLA1 exons 5’ | ccagctttagccttttttcag | POLA1 intron 36 1/4th | ggtctcagaaatccaaacca |
| POLA1 exons 5’ | taaacacctgtgaagtcctc | POLA1 intron 36 1/4th | agaagaggacaggaggtgg g |
| POLA1 exons 5’ | cctgaaccagcttcgaatac | POLA1 intron 36 1/4th | cagtcagctgagtgtcaagg |
| POLA1 exons 5’ | caatccagtcatcatcctgg | POLA1 intron 36 1/4th | taaagttgcggaaggctgct |
| POLA1 exons 5’ | ggcatcatcttcaaggtcat | POLA1 intron 36 1/4th | cgcttggaggacgtgcaaaa |
| POLA1 exons 5’ | tcttgtctttattgcgtgct | POLA1 intron 36 1/4th | aaaccctctactactaccat |
| POLA1 exons 5’ | cactgcgagcttctttacat | POLA1 intron 36 1/4th | acaggacacaggaatatcct |
| POLA1 exons 5’ | tctgcagttttctttccagc | POLA1 intron 36 1/4th | aagagatgagcaccccaaga |
| POLA1 exons 5’ | accatccttggacaagtcta | POLA1 intron 36 1/4th | gacaccccaggtaagatgag |
| POLA1 exons 5’ | cctgtagaatgtcacctagc | POLA1 intron 36 1/4th | cagtcaagctatttctcctg |
| POLA1 exons 5’ | tcagtatcattacaggtggt | POLA1 intron 36 1/4th | actattttataactccagct |

| | | | |
|---|---|---|---|
| POLA1 exons 5’ | tgaagctccaatggatcttt | POLA1 intron 36 $1/4^{th}$ | tccttctaggcaaaaagggg |
| POLA1 exons 5’ | tgtgcacagagaaaggattc | POLA1 intron 36 $1/4^{th}$ | tgggtaccaacaaggtgcag |
| POLA1 exons 5’ | gggaagcaatttttcctgaa | POLA1 intron 36 $1/4^{th}$ | ggatggagtcagaaagtgct |
| POLA1 exons 5’ | agttaatggaggctcctttc | POLA1 intron 36 $1/4^{th}$ | atgacacagttgaaggacat |
| POLA1 exons 5’ | cagcacgtttaagaggaaca | POLA1 intron 36 $1/4^{th}$ | ggtatgggctaaatgtttga |
| POLA1 exons 5’ | tctcgacctgtacatcatcg | POLA1 intron 36 $1/4^{th}$ | aagtccatgtggaatgagga |
| POLA1 exons 5’ | tgactcctgctcttcttctg | POLA1 intron 36 $1/4^{th}$ | gtagacagattaaggtggga |
| POLA1 exons 5’ | ggctcatcaaagtcaccatc | POLA1 intron 36 $1/4^{th}$ | cagcccaactctattagtaa |
| POLA1 exons 5’ | agccataggctccaggtcca | POLA1 intron 36 $1/4^{th}$ | atcttctggagatacatgat |
| POLA1 exons 5’ | tctctttgtcccaagccttg | POLA1 intron 36 $1/4^{th}$ | atttctaggctggaatttat |
| POLA1 exons 5’ | ttgtttcacttcctctgctg | POLA1 intron 36 $1/4^{th}$ | gactgagcatcagtttgatt |
| POLA1 exons 5’ | agtaggacacggtccctttc | POLA1 intron 36 $1/4^{th}$ | ctttcaaacttaatcccact |
| POLA1 exons 5’ | catccgggagaaaacttcct | POLA1 intron 36 $1/4^{th}$ | gttgagacagtcctcatttg |
| POLA1 exons 5’ | tgatcaatgtcccaacaaga | POLA1 intron 36 $1/4^{th}$ | aattctggagggcaaggagt |
| POLA1 exons 5’ | tgcactgagaaactgctatc | POLA1 intron 36 $1/4^{th}$ | gggaaaacaaccagctgcta |
| POLA1 exons 5’ | gactggaatccacttgaact | POLA1 intron 36 $1/4^{th}$ | catctggagcagagagctaa |
| POLA1 exons 5’ | gccccttttaccaatgggag | POLA1 intron 36 $1/4^{th}$ | taaaggctagccaaccaaat |
| POLA1 exons 5’ | agtggaatacttgttcctca | POLA1 intron 36 $1/4^{th}$ | cctccagacatttcaagata |
| POLA1 exons 5’ | ggttggttgtactgatcctc | POLA1 intron 36 $1/4^{th}$ | ttcacgagagctttctggag |
| POLA1 exons 5’ | ctcggctgattcaatccaaa | POLA1 intron 36 $1/4^{th}$ | tctgaaagtctcttagcggt |
| POLA1 exons 5’ | atgacacaacagctcacatg | POLA1 intron 36 $1/4^{th}$ | gctgctcaactgaaagtcac |
| POLA1 exons 5’ | agcgttcgctcgatattttt | POLA1 intron 36 $1/4^{th}$ | attgccattaatacgtgtgc |
| POLA1 exons 5’ | gtttctttccccgtatttag | POLA1 intron 36 $1/4^{th}$ | ccagcctattagtagtattg |
| POLA1 exons 5’ | gcatagttcttttccactgg | POLA1 intron 36 $1/4^{th}$ | cttgtgggctatttttctcc |
| POLA1 exons 5’ | ctggaacatcaggtatctca | POLA1 intron 36 $1/4^{th}$ | tcaagacaggtatctccaga |
| POLA1 exons 5’ | aatcttgaggaagctgtggc | POLA1 intron 36 $1/4^{th}$ | agagtatatcttgcaggtta |
| POLA1 exons 5’ | agatgtgttggtcccaaata | POLA1 intron 36 $1/4^{th}$ | tatcgtgcaatgattactct |
| POLA1 intron 36 3’ | tatcccatagccagtcaata | POLA1 intron 36 $1/4^{th}$ | ttttctaactcgctcagtct |
| POLA1 intron 36 3’ | tgctcttcctcctaaaagac | POLA1 intron 36 $3/4^{th}$ | ttgaactgtgggtatcatct |
| POLA1 intron 36 3’ | agaatgacaaccagcgtcac | POLA1 intron 36 $3/4^{th}$ | taaatcattagccttgtgca |
| POLA1 intron 36 3’ | ctatttacgctgccacacag | POLA1 intron 36 $3/4^{th}$ | catctttagttattcatgca |
| POLA1 intron 36 3’ | accaggttggttagaaaggg | POLA1 intron 36 $3/4^{th}$ | atctctaacaaagttgggga |
| POLA1 intron 36 3’ | ctctgctgatgggatattga | POLA1 intron 36 $3/4^{th}$ | gacaaccgatccattaagga |
| POLA1 intron 36 3’ | atagctcctgatgtgtttac | POLA1 intron 36 $3/4^{th}$ | gagctgtaagttggtgaagt |
| POLA1 intron 36 3’ | aagagctgcggtaatacgga | POLA1 intron 36 $3/4^{th}$ | aggcattttggcaagtttta |
| POLA1 intron 36 3’ | catctgatgactggggagag | POLA1 intron 36 $3/4^{th}$ | aattttttctcccttcagtc |
| POLA1 intron 36 3’ | ccgtgcatgtgaatgacaca | POLA1 intron 36 $3/4^{th}$ | cttaatactagttctctgct |
| POLA1 intron 36 3’ | aaacaacaatggccccaacc | POLA1 intron 36 $3/4^{th}$ | gcggctaatttacattttct |
| POLA1 intron 36 3’ | ctaatgatctgagtctggga | POLA1 intron 36 $3/4^{th}$ | cacttgccttctctaatact |
| POLA1 intron 36 3’ | tacattcgcatttgctctcc | POLA1 intron 36 $3/4^{th}$ | acctgtggtaaaaatggctt |
| POLA1 intron 36 3’ | gtctacaaaacacccggaaa | POLA1 intron 36 $3/4^{th}$ | atgactcagtcagcaacaga |
| POLA1 intron 36 3’ | accggtcaggaaggaaacac | POLA1 intron 36 $3/4^{th}$ | attttaggtttcagtgacct |

| | | | |
|---|---|---|---|
| POLA1 intron 36 3’ | gacatcagctgatcctgagg | POLA1 intron 36 3/4th | gttacgcgatgcttacaaca |
| POLA1 intron 36 3’ | acatagaaatttgcaggcct | POLA1 intron 36 3/4th | tgtcttaagtggatcttgga |
| POLA1 intron 36 3’ | ctgccactacatcactactc | POLA1 intron 36 3/4th | ccatctcagtgatgcagaaa |
| POLA1 intron 36 3’ | tgaggagaggctcacttctg | POLA1 intron 36 3/4th | cttcagcacattattgatgg |
| POLA1 intron 36 3’ | taaggtgtctggagaggctg | POLA1 intron 36 3/4th | ttccctgtatagagagctaa |
| POLA1 intron 36 3’ | atgacgctgtgttcaggaac | POLA1 intron 36 3/4th | acttctgtacattttgctgg |
| POLA1 intron 36 3’ | gaagagagtgatcacgtggg | POLA1 intron 36 3/4th | tgaccagagattccagtgaa |
| POLA1 intron 36 3’ | atgctcaggtctgatctttt | POLA1 intron 36 3/4th | ttagcaacacatgggctgac |
| POLA1 intron 36 3’ | taccagccagagaaaagacc | POLA1 intron 36 3/4th | ccatgggtcaaatcaaacca |
| POLA1 intron 36 3’ | tgctcctttaacccattcat | POLA1 intron 36 3/4th | gtgtaattgcctggattcag |
| POLA1 intron 36 3’ | actgtgtgaactcagggtat | POLA1 intron 36 3/4th | gtggacagtagaacccttaa |
| POLA1 intron 36 3’ | ctgaaagaggcgagaggtga | POLA1 intron 36 3/4th | gacatttctcaagatgagga |
| POLA1 intron 36 3’ | gcctagtaagcattttctag | POLA1 intron 36 3/4th | ttcagatgcaggatatgtgc |
| POLA1 intron 36 3’ | agtctcaaagggagctatgg | POLA1 intron 36 3/4th | ccatttttgtttcttgtgtt |
| POLA1 intron 36 3’ | ctggacgtggcaggaaacac | POLA1 intron 36 3/4th | gtgaaagcactcttcaggca |
| POLA1 intron 36 3’ | cacatgcatgcaactgcttg | POLA1 intron 36 3/4th | cagctgtcccttgattaatg |
| POLA1 intron 36 3’ | agacaaggtgagctctactc | POLA1 intron 36 3/4th | cagacttaattcagcaggcc |
| POLA1 intron 36 3’ | actgatgtgaactgcaggtg | POLA1 intron 36 3/4th | cactcgcagtctgactctta |
| POLA1 intron 36 3’ | ctagctgactcagtgcactg | POLA1 intron 36 3/4th | tcacctaaacacagaggcag |
| POLA1 intron 36 3’ | ttcagtattccaccagagaa | POLA1 intron 36 3/4th | ttacacttgctaaagtggcc |
| POLA1 intron 36 5’ | cagtaggtctggcttttata | POLA1 intron 36 3/4th | tctcacctgagtctatacat |
| POLA1 intron 36 5’ | ctgcttctgagacagcaatg | POLA1 intron 36 3/4th | cccaaactcaaggggcaaaa |
| POLA1 intron 36 5’ | gccttgctttaatagaggac | POLA1 intron 36 3/4th | agatctggaggtcatttctc |
| POLA1 intron 36 5’ | agagtccttttttgtactgtt | POLA1 intron 36 3/4th | tcctccacagtgaatgatat |
| POLA1 intron 36 5’ | gttgcctatgcttactttat | POLA1 intron 36 3/4th | atccctaaggggaaattacc |
| POLA1 intron 36 5’ | agcgaacagcagagacaca c | POLA1 intron 36 3/4th | tctccctattaatgcataca |
| POLA1 intron 36 5’ | gcctatatactctcttagga | POLA1 intron 36 3/4th | atttgtgactctctctctct |
| POLA1 intron 36 5’ | tgcaacagctgatcatacgg | POLA1 intron 36 3/4th | aggcatagacacttaccata |
| POLA1 intron 36 5’ | ccgcaagtcttgtataagga | POLA1 intron 36 3/4th | ttttcctgacgagttctaca |
| POLA1 intron 36 5’ | cttatgcggggaaatctagc | POLA1 intron 36 3/4th | aaatgcatagtcgtcacctg |
| POLA1 intron 36 5’ | gcctatcttgtggatggaaa | POLA1 intron 36 3/4th | ggtaagagtcggatctgtta |
| POLA1 intron 36 5’ | tctccaaggacagcaaaacc | POLA1 intron 36 3/4th | atgttgttaactggtcgtct |
| POLA1 intron 36 5’ | atgctgtggagatagctcaa | POLA1 intron 36 3/4th | tcaaaaaccatcactgcctt |
| POLA1 intron 36 5’ | atatgggctagacattgacc | POLA1 intron 36 3/4th | ttcagagagaaactcgaggt |
| POLA1 intron 36 5’ | actgtctttaggaaaggtcc | POLA1 intron 36 3/4th | ctaggggtttctacagtgtg |
| POLA1 intron 36 5’ | gccgtatagggaagaggaat | POLA1 intron 36 3/4th | aattctgtgcctgacagagc |
| POLA1 intron 36 5’ | aaagactgattctgcctctc | POLA1 intron 36 3/4th | attaatttttcttccctggg |
| POLA1 intron 36 5’ | agttgccagaccttttgaaa | POLA1 intron 36 3/4th | aagagaacacagccttctcc |
| POLA1 intron 36 5’ | aaatctccagtgactttcca | POLA1 intron 36 3/4th | gctcttgctgctaaatatca |
| POLA1 intron 36 5’ | cagtgtcctctaagaggatg | POLA1 intron 36 3/4th | aagaaggaaacatccctctc |
| POLA1 intron 36 5’ | gtgggtgtaataacgtgcac | POLA1 intron 36 3/4th | gtgtttattgtaactactgc |
| POLA1 intron 36 5’ | gatcctgtcaatacttttcc | POLA1 intron 36 3/4th | gagtcctctgagatttggaa |

| | | | |
|---|---|---|---|
| POLA1 intron 36 5’ | tcactcattcccaaactgta | POLA1 intron 35 3’ | acagccaagtcatttgtaca |
| POLA1 intron 36 5’ | gcctggagcatgtaaatttc | POLA1 intron 35 3’ | ctgccttggaatgacatgta |
| POLA1 intron 36 5’ | ataccctaataccacttagc | POLA1 intron 35 3’ | aacatgttaccagtttctcc |
| POLA1 intron 36 5’ | agttttggtttatatcccac | POLA1 intron 35 3’ | agccctcagaacttttcaaa |
| POLA1 intron 36 5’ | atgtgttaggaaacctctgc | POLA1 intron 35 3’ | gtcctaccaaggtgaaacaa |
| POLA1 intron 36 5’ | aggagagggcttatcagatg | POLA1 intron 35 3’ | gctttggtgttttctttgag |
| POLA1 intron 36 5’ | cctatgaactatgaccttgc | POLA1 intron 35 3’ | ttgcgagctggggaacaatg |
| POLA1 intron 36 5’ | gttgctaggtactgacagta | POLA1 intron 35 3’ | aggggttagacacttgggaa |
| POLA1 intron 36 5’ | ggggttaatccacattatgg | POLA1 intron 35 3’ | aaactttctttccatcttcc |
| POLA1 intron 36 5’ | tatgccatctgtgtacattc | POLA1 intron 35 3’ | ggggagtcacattcaaatgc |
| POLA1 intron 36 5’ | cctatgttcctgtaaaaccg | POLA1 intron 35 3’ | cctgaggttatttagagtca |
| POLA1 intron 36 5’ | aggaatgtcactcaacacct | POLA1 intron 35 3’ | gtaggagacgtgtctgaact |
| POLA1 intron 36 5’ | taatttcaccctattggtgg | POLA1 intron 35 3’ | cttgccaagttccaattcaa |
| POLA1 intron 36 middle | ctgtggaacttaatggccat | POLA1 intron 35 3’ | cacccatgtcaaaacactcg |
| POLA1 intron 36 middle | agcaggtggtagcattatag | POLA1 intron 35 3’ | tatgatctctgagaagtacc |
| POLA1 intron 36 middle | ctattcattccacacctatt | POLA1 intron 35 3’ | tggtaatcctaagtgggcaa |
| POLA1 intron 36 middle | tcctatgaatctgcagcata | POLA1 intron 35 3’ | aacatcttgactatttgctc |
| POLA1 intron 36 middle | gtgagatttcatggggttac | POLA1 intron 35 3’ | gattggctcacagactcaca |
| POLA1 intron 36 middle | ttgggaccattcttatatag | POLA1 intron 35 3’ | ttgtggtgacaatccattgc |
| POLA1 intron 36 middle | acccacagttcaaagaacta | POLA1 intron 35 3’ | atattgctataccatgagcc |
| POLA1 intron 36 middle | cacagattctactattcctt | POLA1 intron 35 3’ | tgctatgtgcttcctaactg |
| POLA1 intron 36 middle | ctttagaactgaatgcctcc | POLA1 intron 35 3’ | atttgcactgttaattggga |
| POLA1 intron 36 middle | atcagacatgccattgagtt | POLA1 intron 35 3’ | actgcaaattcctatgcact |
| POLA1 intron 36 middle | gctctttcttcatcacgatg | POLA1 intron 35 3’ | cctgctgtcaaaaacctgat |
| POLA1 intron 36 middle | tgagcaagagttggggcaag | POLA1 intron 35 3’ | attactacttgcgaagctgc |
| POLA1 intron 36 middle | gatcacttacaatcctgtgc | POLA1 intron 35 3’ | ataaatcctgccacttcgat |
| POLA1 intron 36 middle | ggctgtaacccctgaaagag | POLA1 intron 35 3’ | tagcatatacacctttcata |
| POLA1 intron 36 middle | ccaaacctatgggttctcta | POLA1 intron 35 3’ | ctaatggtttgtccctctta |
| POLA1 intron 36 middle | aggaacccaggttttctgac | POLA1 intron 35 3’ | gggtaagaagggcagaatca |
| POLA1 intron 36 middle | ctcacagcacagtaatcttc | POLA1 intron 35 3’ | cactctcagcagagtgtatc |

| | | | |
|---|---|---|---|
| POLA1 intron 36 middle | gtccctattttaagactgga | POLA1 intron 35 3’ | tgcctctgcatttaagaagt |
| POLA1 intron 36 middle | actctttaaaggacaggcat | POLA1 intron 35 3’ | tgtgtatcaacccttttcag |
| POLA1 intron 36 middle | gtggcatttactggatttca | POLA1 intron 35 3’ | ctgtgtgtacatatttttct |
| POLA1 intron 36 middle | tgggattttgctgacatagt | POLA1 intron 35 3’ | cctgactagttgccaaattt |
| POLA1 intron 36 middle | gcagggagttattagtttgg | POLA1 intron 35 3’ | acgatggtagctatttgcag |
| POLA1 intron 36 middle | cttctacgctgccaaaacag | POLA1 intron 35 3’ | ttcattgtgatggcaaccat |
| POLA1 intron 36 middle | ctccttcaaatttcacaact | POLA1 intron 35 5’ | ctttagcattatgagactgt |
| POLA1 intron 36 middle | cttaagagtaacggcagcac | POLA1 intron 35 5’ | ctctaccctcttctacatgg |
| POLA1 intron 36 middle | acttgactcaattagcacct | POLA1 intron 35 5’ | tctacttctcaaagcaagca |
| POLA1 intron 36 middle | gtgcctgcattagatcataa | POLA1 intron 35 5’ | cttcatagggcaaatactca |
| POLA1 intron 36 middle | tgacaccctgacatctgatc | POLA1 intron 35 5’ | tatgcatatggtagtagggc |
| POLA1 intron 36 middle | aatgcctttgggtagagctt | POLA1 intron 35 5’ | gaaagcatgtatctttctga |
| POLA1 intron 36 middle | ctagctagaaggtgtgtctg | POLA1 intron 35 5’ | gtattgctcatgacatgtga |
| POLA1 intron 36 middle | gtctgtgaattggtggatct | POLA1 intron 35 5’ | gaattagtcagtttcccttt |
| POLA1 intron 36 middle | agcatgaagtggcatatgac | POLA1 intron 35 5’ | gtaattatgtagtggcaacc |
| POLA1 intron 36 middle | gcgcaaggcagtaagctaaa | POLA1 intron 35 5’ | aatggaatggccatgtcttc |
| POLA1 intron 36 middle | ttcaatacttcttgtgtgct | POLA1 intron 35 5’ | cccaaagacgatagcagttt |
| POLA1 intron 36 middle | tttatttggtgtcagctacc | POLA1 intron 35 5’ | ctggctcttacaaatgggat |
| POLA1 intron 36 middle | cttgaaattacagattccct | POLA1 intron 35 5’ | gggtacagaattgggaggat |
| POLA1 intron 36 middle | tgcctttcaaaatacgcagt | POLA1 intron 35 5’ | accccacaaaataaaccatt |
| POLA1 intron 36 middle | aggtgaatgtgactggtgtc | POLA1 intron 35 5’ | caattaatccagcagagggg |
| POLA1 intron 36 middle | tctgtgaggaactactcaga | POLA1 intron 35 5’ | ctactcatctatgcagctac |
| POLA1 intron 36 middle | aaccatatgtttttccccat | POLA1 intron 35 5’ | tacacctcaggtgtgtatat |
| POLA1 intron 36 middle | gcctatggaattatgagact | POLA1 intron 35 5’ | ctttctagttgggagtcatt |
| POLA1 intron 36 middle | ggctgaggacagctacaaat | POLA1 intron 35 5’ | agaagtgtttccagatgcca |

| | | | |
|---|---|---|---|
| POLA1 intron 36 middle | aggtttcttaccatcttcat | POLA1 intron 35 5’ | gggaaggcatcttaattacc |
| POLA1 intron 36 middle | ggtatagatagcttaccagc | POLA1 intron 35 5’ | gatgctcaaaggggtcaaca |
| POLA1 intron 36 middle | aattgctgcagtccattcac | POLA1 intron 35 5’ | ggaaaatgctgtttgggtcg |
| POLA1 intron 36 middle | acaaggttaactaggttcgt | POLA1 intron 35 5’ | gaacctgcaaacttctcact |
| POLA1 intron 36 middle | agtgatctgatataggggga | POLA1 intron 35 5’ | ccttcttctcactaaaaacc |
| POLA1 intron 36 middle | ccctctactgggaattttaa | POLA1 intron 35 5’ | aaacaacaggttcccaagtc |
| POLA1 intron 36 middle | gtttgacatccagtcagatt | | |
| POLA1 intron 36 middle | acagtgtacaacggatacca | | |
| POLA1 intron 36 middle | ctagcacatactgattctgc | | |
| POLA1 intron 36 middle | tctggtaggcttttagtttg | | |
| POLA1 intron 36 middle | ctgatccttcacttaccaag | | |
| POLA1 intron 36 middle | aagaagccagatttgggctg | | |
| POLA1 intron 36 middle | taactctatctggagaccac | | |
| POLA1 intron 36 tiling | cctctctttgcttccaaaat | | |

**Table S 2: Probe combinations used**

| Probes Set# | Experiment | Combination of probes/antibodies and dyes used (from Table S2) |
|---|---|---|
| 1 | MDN1 middle alternating probes | MDN1 middle odd – Cy5 |
| | | MDN1 middle even – Cy3 |
| 2 | MDN1 5’-middle-3’ | MDN1 5’+ MDN1 5’ additional – Dy488 |
| | | MDN1 middle – Cy5 |
| | | MDN 3’ – Dy550 |
| 3 | POLA1 intron 36-exon | POLA1 5’ exon Cy3 |
| | | POLA1 intron 36 5’ Cy5 |
| 4 | POLA1 intron 35-exon | POLA1 5’ exon Cy3 |
| | | POLA1 intron 35 5’ Cy5 |
| 5 | POLA1 exons 5’-3’ intron 36 middle | POLA1 5’ exons -Cy3 |
| | | POLA1 3’ exons- Cy5 |
| | | POLA1 intron 36 middle Atto488 |
| 6 | POLA1 intron 36 5’-middle-3’ | POLA1 intron 36 5’ Cy5 |
| | | POLA1 intron 36 middle Atto488 |
| | | POLA1 intron 36 3’ Cy3 |
| 7 | POLA1 intron 36 tiling middle 3’ | POLA1 intron 36 tiling Cy5 |
| | | POLA1 intron 36 middle Atto488 |
| | | POLA1 intron 36 3’ Cy3 |
| 8 | POLA1 intron 36 5’ 1/4th middle | POLA1 intron 36 5’ Cy5 |
| | | POLA1 intron 36 1/4th Cy3 |
| | | POLA1 intron 36 middle Atto488 |
| 9 | POLA1 intron 36 1/4th middle-3/4th | POLA1 intron 36 1/4th Cy3 |
| | | POLA1 intron 36 middle Atto488 |
| | | POLA1 intron 36 3/4th Cy5 |
| 10 | POLA1 intron 36 middle-3/4th-3’ | POLA1 intron 36 middle Atto488 |
| | | POLA1 intron 36 3/4th Cy5 |
| | | POLA1 intron 36 3’ Cy3 |

| | | |
|---|---|---|
| 11 | POLA1 intron 36 5’-1/4th-3’ | POLA1 intron 36 5’ Cy5 |
| | | POLA1 intron 36 1/4th Atto488 |
| | | POLA1 intron 36 3’ Cy3 |
| 12 | POLA1 intron 36 5’-3/4th-3’ | POLA1 intron 36 5’ Cy5 |
| | | POLA1 intron 36 3/4th Atto488 |
| | | POLA1 intron 36 3’ Cy3 |
| 13 | POLA1 intron 35 5’-1/3rd-3’ | POLA1 intron 35 5’ Cy5 |
| | | POLA1 intron 35 1/3rd Atto488 |
| | | POLA1 intron 35 3’ Cy550 |
| 14 | POLA1 intron 35 5’-2/3rd-3’ | POLA1 intron 35 5’ Cy5 |
| | | POLA1 intron 35 2/3rd Atto488 |
| | | POLA1 intron 35 3’ Cy550 |
| 15 | AHNAK 5’-middle-3’ | AHNAK 5’ – Dy488 |
| | | AHNAK middle – Cy5 |
| | | AHNAK 3’ – Cy3 |
